# Inhibition of cytosolic DNA sensing and transposon activity safeguards pluripotency

**DOI:** 10.1101/2025.03.26.645264

**Authors:** Ferran Garcia-Llagostera, Audrey LK Putman, Alexandros Choromidis, Bryony J Leeke, Klaudia Stanik, Antonia Ramos-Guzmán, Benjamin Moyon, Jesus Gil, Alexis R. Barr, Michelle Percharde

## Abstract

Transposable elements (TEs) are mobile DNA sequences that make up a sizeable fraction of mammalian genomes yet are often tightly repressed by transcriptional and epigenetic mechanisms. During early development, epigenetic reprogramming selectively loosens TE repression, and TE transcription actively contributes to embryogenesis. This raises the question: how can embryos and embryonic stem cells (ESCs) tolerate TE expression without incurring widespread inflammation or DNA damage? Here, we reveal multiple mechanisms that prevent innate immune activation by TE-derived cytosolic DNA, including reduced cGAS/STING expression and signalling, dampening of Type I interferon responses by pluripotency factors, and post-transcriptional restriction of retrotransposition. These layers of protection are essential, as experimental perturbation triggers loss of ESC self-renewal and pluripotency. Our data explain how early development can be permissive to TE expression while safeguarding against harmful effects of TE activity.

## Introduction

A significant portion of mammalian genomes is comprised of transposable elements (TEs): DNA sequences that are either historically – or still currently – mobile. Class I TEs are retrotransposons that mobilise via an RNA intermediate that is reverse transcribed back into cDNA and inserted into a new location. In human, LINE1 is the only active and autonomous retrotransposon family, with approximately 100 retrotransposition-competent copies in human and around 2000 in mouse (Goodier et al. 2001; Brouha et al. 2003; Kazazian and Moran 2017). However, some TEs belonging to both LINE1 and LTR families maintain enzymatic activity and can generate complementary DNA (cDNA) via reverse transcription (Kassiotis and Stoye 2016).

TEs are tightly repressed in most cell types, primarily at the transcriptional and epigenetic level (Bestor and Bourc’his 2004; Almeida et al. 2022), as well as via small RNA-based silencing in specific cell types (Tam et al. 2008; Castaneda et al. 2011; Ozata et al. 2019). This is thought to be because both TE expression and the TE replication cycle can perturb cellular homeostasis by multiple mechanisms. TE-derived RNA can play roles in the nucleus, mediating chromatin or gene regulatory effects (Percharde et al. 2018; Della Valle et al. 2022; Marasca et al. 2022). A well-known consequence of TE activity is DNA damage or insertional mutagenesis; this has been shown to drive tumorigenesis (Iskow et al. 2010; Lee et al. 2012; Shukla et al. 2013; Burns 2017), to contribute to infertility (Malki et al. 2014; Newkirk et al. 2017; Zoch et al. 2024) as well as to cause heritable disorders (Kazazian et al. 1988; Chen et al. 2005). Recently, a growing body of evidence points to TE-driven activation of the innate immune system, which causes inflammation and contributes to senescence (De Cecco et al. 2019; Simon et al. 2019; Liu et al. 2023; Zhang et al. 2023).

The innate immune system is a conserved defence system present in all multicellular organisms. Invading pathogens or threats contain certain molecular features, termed pathogen-associated molecular patterns (PAMPs), which are recognised by innate immune receptors called pattern recognition receptors (PRRs). The activation of PRRs triggers an inflammatory response which stimulates effectors while also helping to initiate an adaptive immune response (Carpenter and O’Neill 2024). Viral-derived double-stranded (dsRNA) or cDNA can both act as PAMPs, activating the RIG-I-MDA5-MAVS pathway or cGAS-STING pathway, respectively. Both pathways converge on the phosphorylation and activation of TBK1 followed by IRF3/7, resulting in transcription of Type I interferons (Type-I IFNs) and inflammatory cytokines. IFNs diffuse out of the cell, activating JAK/STAT signalling in the same and neighbouring cells and amplifying an immune response. Critically, it was recently shown that TE-derived dsRNA or cytosolic DNA produced from TE reverse transcriptase can trigger an IFN-I response (Gao et al. 2013; Chiappinelli et al. 2015; Kassiotis and Stoye 2016; Zhao et al. 2021). Innate immune activity and inflammation have been directly linked to the pathological effects of TE reactivation in multiple contexts (Thomas et al. 2017; De Cecco et al. 2019; Simon et al. 2019; Zhao et al. 2021; Scopa et al. 2023).

In contrast to their strict repression in somatic cells, many TEs are transcriptionally upregulated during the global epigenetic reprogramming that occurs in early development (Macfarlan et al. 2011; Seisenberger et al. 2012; Ohno et al. 2013; Jachowicz et al. 2017; Percharde et al. 2018). Moreover, LINE1 and LTR elements can act as important developmental regulators of gene expression (Macfarlan et al. 2012; Grow et al. 2015; Percharde et al. 2018; Modzelewski et al. 2021). Indeed, to have been propagated during evolution, TEs must also have occasionally mobilized and been transmitted through the germline (Kano et al. 2009; Ewing and Kazazian 2010; Richardson et al. 2017). Thus, there must be specific mechanisms that prevent the sensing of expressed TEs by PRRs as well as mechanisms that limit TE retrotransposition activity to low levels in early development. However, such mechanisms are unknown or poorly understood.

Focusing on LINE1, we asked how TE expression can be tolerated without pathology in early development. We investigated cGAS/STING that detects cytosolic DNA and reveal that this pathway is repressed at multiple levels in early development, rendering ESCs insensitive to potential TE activity. Moreover, we reveal that LINE1 is highly expressed in ESCs yet its retrotransposition activity is heavily repressed. Rescuing the sensing of either TE expression or activity by constitutive IRF3 activation severely compromises self-renewal and pluripotency. At the same time, our data suggest that pluripotency factors limit the IFN-I response in ESCs. We propose that repression of the pathways that sense TE expression and activity permits the expression of these elements, while simultaneously guarding against TE-driven pathology.

## Results

### ESCs do not respond to cytosolic DNA

ESCs and embryos show transcriptional activation of several TE subfamilies yet seem to avoid pathological consequences of their de-repression. This is unlike somatic or cancer cells, where TE expression or activity can induce an innate immune response (Kassiotis and Stoye 2016). We thus set out to investigate the relationship between TE sensing and innate immune activation in pluripotent cells. High LINE1 RNA expression has been previously documented in ESCs and pre-implantation embryos, which can be translated into two proteins, ORF1p and ORF2p; both are essential for LINE1 retrotransposition activity (Moran et al. 1996). While ORF2p is undetectable in most cells and tissues (Ardeljan et al. 2020b; Nielsen et al. 2025), ORF1p is an excellent marker for LINE1 expression and activity (Rodic et al. 2014). We profiled ORF1p in pluripotent ESCs and upon undirected differentiation and found that ORF1p is abundant in undifferentiated ESCs (**Fig.1A**). Moreover, it is rapidly downregulated upon exit from pluripotency (**Fig.1B**).

**Figure 1.**
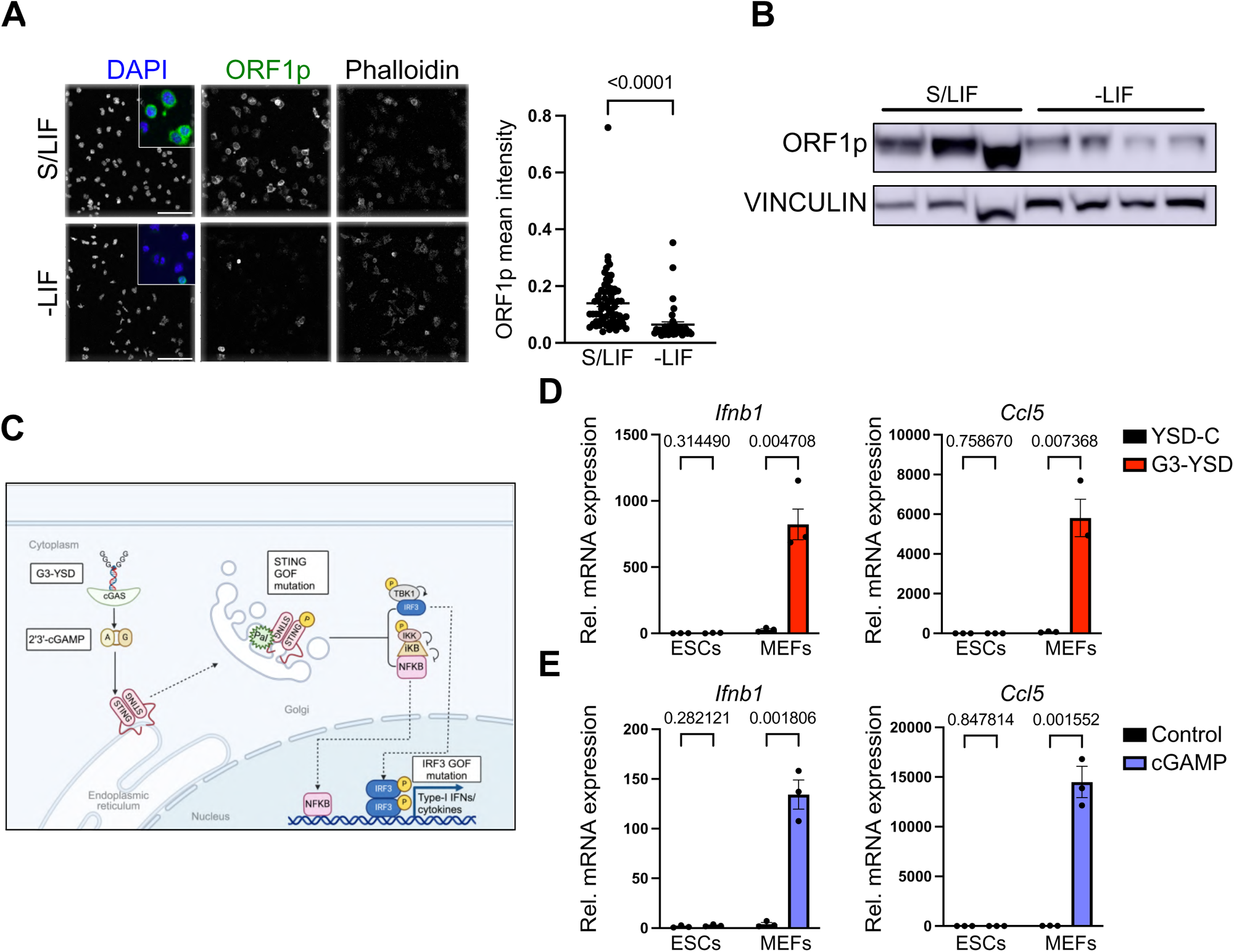
Sensing of TE activity by cGAS/STING is abrogated in ESCs. a) Immunofluorescence and quantification of LINE1 ORF1p levels in ESCs in pluripotent conditions, or induced to differentiate by LIF withdrawal. Mean cytoplasmic signals per cell are plotted in each condition, with P values from Welch’s t-test. Data representative of 3 experiments. Scale bar, 100μm. b) Western blotting of ORF1p levels in experiments performed as in a). Vinculin is shown as a loading control. Samples from 3-4 experiments are shown. c) Diagram of cGAS/STING signaling. Cytoplasmic DNA activates cGAS to produce cGAMP, which in turns activates STING and induces its transit from the ER to Golgi. There, phosphorylated STING phosphorylates and activates TBK1, ultimately inducing phosphorylation and activation of the transcription factor, IRF3. IRF3 binds and activates transcription of Type-I interferons and other cytokines. Made with Biorender. d) Induction of the indicated genes in ESCs and MEFs 6 h after transfection of control (YSD-C) or STING-activating (G3-YSD) DNA ligands. e) As in d) but 6 h after control (mock) transfection or transfection of cGAMP. Data in d-e are mean +/- s.e.m, n=3 individual wells, representative of at least 2 experiments. P values in d and e, two-tailed Student’s t-test with Holm-Šídák correction.

Given the abundance of LINE1 in ESCs, we first interrogated the cGAS/STING pathway, which detects cytoplasmic DNA such as that generated by LINE1 activity (**Fig.1C**). We transfected ESCs and a somatic cell model, mouse embryonic fibroblasts (MEFs), with palindromic Y-shaped 16-mer DNA that is a potent cGAS agonist (G3-YSD, (Herzner et al. 2015)). G3-YSD, but not a control DNA (YSD-C) induces a rapid interferon response in MEFs (**Fig.S1**). In contrast to MEFs, we found that G3-YSD transfection has no effect in ESCs on either *Ifnb1* expression or the interferon stimulated gene (ISG), *Ccl5* (**Fig.1D, S1**). STING is the adaptor for cGAS, but also other PRRs such as DDX41 and IFI16 (Paludan and Bowie 2013). We activated STING directly with its ligand, 2’3-cGAMP, which also elicits no induction of *Ifnb1* or *Ccl5* in ESCs, despite potent activation in MEFs (**Fig.1E**). Thus, ESCs are insensitive to cytosolic DNA and cGAS/STING activation.

### cGAS/STING expression and activity are attenuated in ESCs

To investigate the mechanism behind the lack of an interferon response in ESCs, we profiled the RNA expression of known factors involved the cGAS/STING pathway. We found that all components examined are expressed in ESCs, albeit at 2-5 fold lower expression than in MEFs (**Fig.2A, S2A**). At the protein level, both TBK1 and IRF3 are detected in ESCs yet not phosphorylated upon cytoplasmic DNA stimulation, confirming a lack of cGAS/STING signalling (**Fig.2B**). Strikingly, cGAS and STING protein are absent in ESCs, and not rescued upon early passive differentiation (**Fig.2C, S2B**), potentially explaining the insensitivity of ESCs to cytosolic DNA.

**Figure 2.**
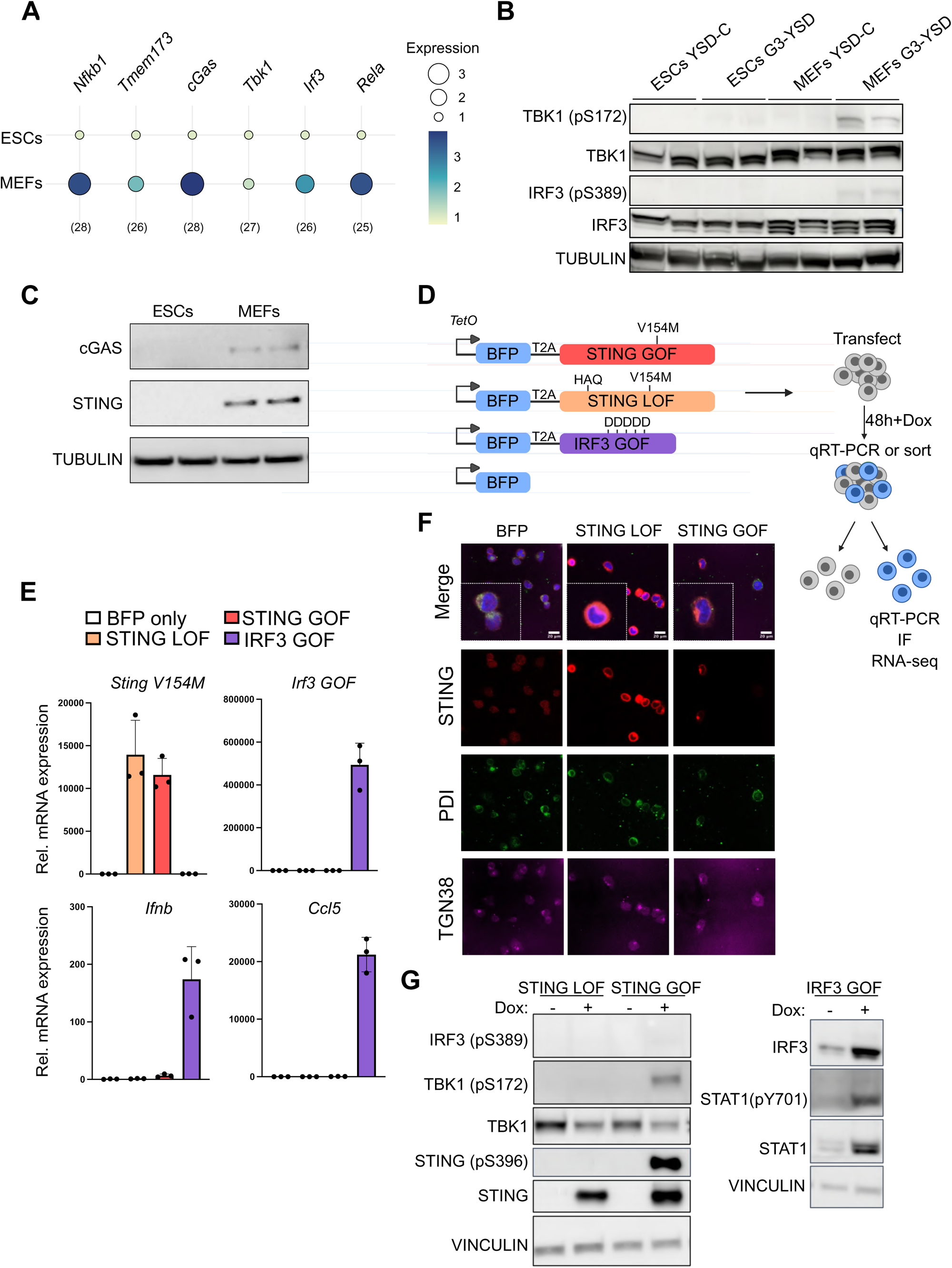
STING is unable to induce an innate immune response in ESCs. a) Bubble plot of mean RNA expression of key factors involved in DNA sensing in ESCs vs MEFs determined by RT-qPCR (n=3), representative of 3 experiments. b) Western blot of the indicated factors in ESCs and MEFs, showing a lack of TBK1 or IRF3 phosphorylation in ESCs. Samples from 2 individual wells are shown. c) Western blot in ESCs and MEFs showing the absence of cGAS and STING protein in ESCs, representative of 3 experiments. TUBULIN is shown as a loading control. d) Diagram of pCW57.1-based Doxycycline(Dox)-inducible constructs used in subsequent experiments. ESCs were transfected with the indicated constructs in the presence of Dox, and BFP-expressing cells sorted by FACS 48 h later for downstream analyses. e) RT-qPCR analysis of the indicated genes in ESCs following transfection and sorting. *Sting V154M* and *Irf3 GOF* primers are mutation specific. Data show mean +/- s.e.m of 3 individual wells. f) Immunofluorescence experiments localizing STING LOF/GOF proteins to the ER (PD1) or Golgi (TGN38) in ESCs. BFP transfected cells are included as a negative control. g) Western blots of the indicated proteins in stable LOF/GOF ESC lines, 24 h after Dox addition. All data in the figure are representative of at least 2 independent experiments unless indicated otherwise.

STING-associated vasculopathy with onset in infancy (SAVI) is an interferonopathy driven by a gain of function (GOF) point mutation in STING (V155M) that renders it constitutively active (Cerboni et al. 2017). To rescue cGAS/STING signalling, we transfected ESCs with a construct expressing BFP and the analogous mouse STING GOF variant (V154M, STING GOF (Bouis et al. 2019)) (**Fig.2D**). As controls, ESCs were transfected with BFP alone or a STING loss of function (LOF) mutant (V154M HAQ; (Patel et al. 2017)), then sorted for BFP fluorescence and analysed by RT-qPCR (**Fig.2D**). Despite inducing *IFNB1* expression when transfected into 293T cells (**Fig.S2C**), STING GOF surprisingly has no effect in ESCs (**Fig.2E**). We verified that STING GOF localises to the Golgi as expected, while STING LOF remains mainly in the ER (**Fig.2F**). To permit stable over-expression in ESCs, cell lines were generated containing Doxycycline (Dox)-inducible proteins expressed from the *Col1a1* locus (KH2 ESCs, (Beard et al. 2006)) (**Fig.S2D-F**). Western blot analysis upon Dox treatment revealed that although STING and TBK1 are phosphorylated by activated STING, phospho-IRF3 is absent (**Fig.2G**). To confirm that IRF3 activation is the limiting step in ESCs, we generated a constitutively active IRF3 construct, which we refer to as IRF3 GOF (**Fig.S2F**)(IRF3-5D, (Lin et al. 1999; Wang et al. 2014a)). Transient transfection of IRF3 GOF leads to a robust induction of *Ifnb1* expression as well as of direct IRF3 target cytokines such as *Ccl5* (**Fig.2E, S2E**). In stable cell lines, IRF3 GOF induction additionally promotes STAT1 phosphorylation – a marker of downstream IFNβ signalling (**Fig.2G**). Taken together, our data reveal that ESCs are insensitive to cytosolic DNA, due to a strict repression of both cGAS/STING expression and downstream signalling.

### Post-transcriptional repression of LINE1 activity is a second line of defence in ESCs

Our results point to a strong mechanism to prevent canonical innate immunity in ESCs, which we hypothesize is at least partly to avoid sensing of TEs, which are highly expressed in development. However, TEs may also cause pathology by DNA damage or insertional mutagenesis. We therefore next asked whether expressed TEs like LINE1 are able to retrotranspose in ESCs, as their high abundance (**Fig.1A-B**) might suggest.

We took advantage of a well-characterised reporter assay to read out the retrotransposition ability of LINE1 in ESCs (L1ORFeus, (An et al. 2006)). We found that even upon correction for differences in transfection efficiency, ESCs exhibit a strong repression of retrotransposition in comparison to 293T cells (**Fig.3A**). To compare our findings with endogenous LINE1 levels in a somatic context, we set up a model of stress-induced senescence in MEFs, in which LINE1 expression and activity has been documented to increase (De Cecco et al. 2019; Simon et al. 2019) (**Fig. S3A-B**). Late-senescent MEFs indeed upregulate LINE1 RNA and ORF1p protein levels, yet this is much lower than endogenous ORF1p in ESCs (**Fig.3B-C, S3C**). Thus, lower LINE1 RNA and/or ORF1p levels do not explain the repression of LINE1 activity in pluripotency. To interrogate the composition of LINE1 complexes in ESCs, we carried out co-immunoprecipitation of endogenous ORF1p in ESCs, first validating its interaction with MOV10 (**Fig.3D** (Goodier et al. 2012)). Profiling ORF1p-associated proteins by IP-mass spectrometry revealed many known interactors of LINE1, as well as novel protein partners (**Fig.3E, S3D**). However, proteins that bind ssDNA or have been associated with LINE1 integrations in other cells (RPA1, PCNA, XRCC5, XRCC6 (Miyoshi et al. 2019; Ardeljan et al. 2020a)) are depleted or not enriched in ESCs (**Fig.3E**). Additionally, we performed staining for the presence of cytosolic DNA – a marker of LINE1 ORF2p activity (Thomas et al. 2017; De Cecco et al. 2019). Despite abundant LINE1 expression, no cytoplasmic ssDNA staining is detected in ESCs, unlike in senescent MEFs (**Fig.3F**). Taken together, our data reveal that the sensing of TEs as well as TE retrotransposition activity is heavily suppressed in pluripotency.

**Figure 3.**
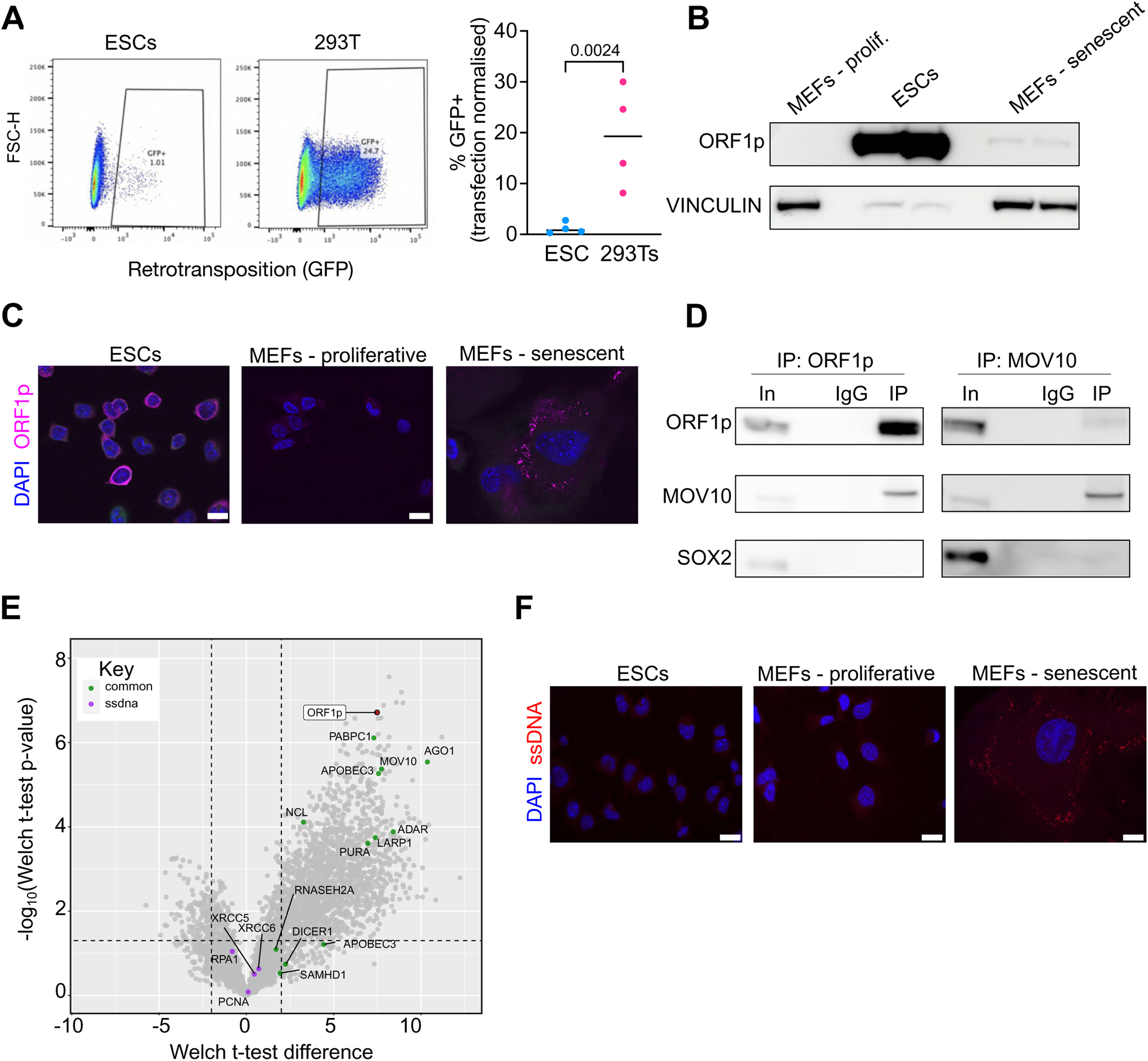
ESCs heavily restrict LINE1 activity. a) Flow cytometry analysis of LINE1 GFP reporter activity in ESCs and 293T cells, carried out 72 h after transfection with L1 ORFeus retrotransposition reporter plasmid (An et al. 2006). To normalize for differing transfection efficiencies, both cell types were co-transfected with 1/10^th^ amount of mCherry-expressing plasmid, and only mCherry-positive cells analyzed. Shown are 4 independent experiments, P value, paired ratio t-test. b) Western blot of LINE1 ORF1p levels in proliferative or senescent MEFs versus ESCs. Vinculin is shown as a loading control; note that much less ESC lysate was loaded than MEF lysate in order to not overwhelm the ORF1p blot. c) Immunofluorescence analysis of ORF1p levels in ESCs versus proliferative or senescent MEFs. d) Co-IP experiments in ESCs confirming successful immunoprecipitation of endogenous ORF1p and validation of its interaction with endogenous MOV10, representative of >3 experiments. e) Volcano plot of ORF1p associated proteins identified by IP-MS in ESCs. Previously reported ORF1p interactors in somatic/cancer cells are also enriched in ESCs (green). ssDNA binding proteins are depleted from ORF1p interactors in ESCs (purple). Dotted lines denote Welch P value = 0.05, and Welch t-test difference = −2 and +2. f) ssDNA staining in proliferative or senescent MEFs versus ESCs, showing positive ssDNA foci only in senescence. All data in figure are representative of at least 2 independent experiments unless otherwise indicated.

### IRF3 activation is incompatible with pluripotency

The attenuation of the cGAS/STING pathway suggested a strong pressure in early development to prevent sensing of TEs or other cytosolic DNA. To understand why, we focused on IRF3 as the downstream effector of the cGAS/STING pathway. Notably, IRF3 phosphorylation is also activated upon sensing of dsRNA by the RIG-I/MDA5/MAVs pathway, which is also non-functional in ESCs for unknown reasons (Chen et al. 2010; Wang et al. 2013; Witteveldt et al. 2019).

To explore the consequences of TE sensing and IRF3 activation, we profiled ESCs by RNA-seq after transfection of STING or IRF3 GOF constructs and sorting for BFP (**Fig.2D, Table S1**). Clustering of samples by PCA revealed that IRF3 GOF-expressing ESCs are distinct from BFP and STING-transfected ESCs (**Fig.4A**). In contrast, only small differences are seen upon STING over-expression, in agreement with its inability to activate IRF3 (**Fig.2E,G**). We confirmed that IRF3 induces the expression of *Ifnb1* as well as many other ISGs, in addition to stimulating interferon-driven gene expression (**Fig.4B, S4A**). A mild upregulation of some ISGs was apparent upon STING GOF, but to much lower levels (**Fig.S4B**) – again consistent with inhibition of signalling downstream of STING in ESCs **(Fig.2G).** Apoptosis-related genes are also upregulated in response to the over-expression of IRF3, but not STING (**Fig.S4C**), as has been described in neutrophils (Andzinski et al. 2015). Strikingly, we found that IRF3 activation causes rapid downregulation of many pluripotency genes, including *Nanog, Pou5f1, Esrrb, Klf2/4/5* and an overall reduction in processes related to self-renewal and development (**Fig.4B-C**). Accordingly, ESC self-renewal, assessed in colony formation assays, is significantly disrupted compared to BFP or STING overexpression (**Fig.4D**). Immunofluorescence analysis in ESCs revealed that IRF3 activation induces the loss of essential pluripotency proteins such as NANOG and SOX2 (**Fig.4E-F**). Using IRF3 GOF stable cell lines (**Fig.S2D-E, S4D**), we confirmed that IRF3 activation induces a more differentiated morphology after only a few days, and concomitant loss of ESC self-renewal capacity (**Fig.4G-I, S4D-E**). These data reveal that IRF3 activation antagonises pluripotency, suggesting that activation of the pathways that would be involved in TE sensing is incompatible with early development.

**Figure 4.**
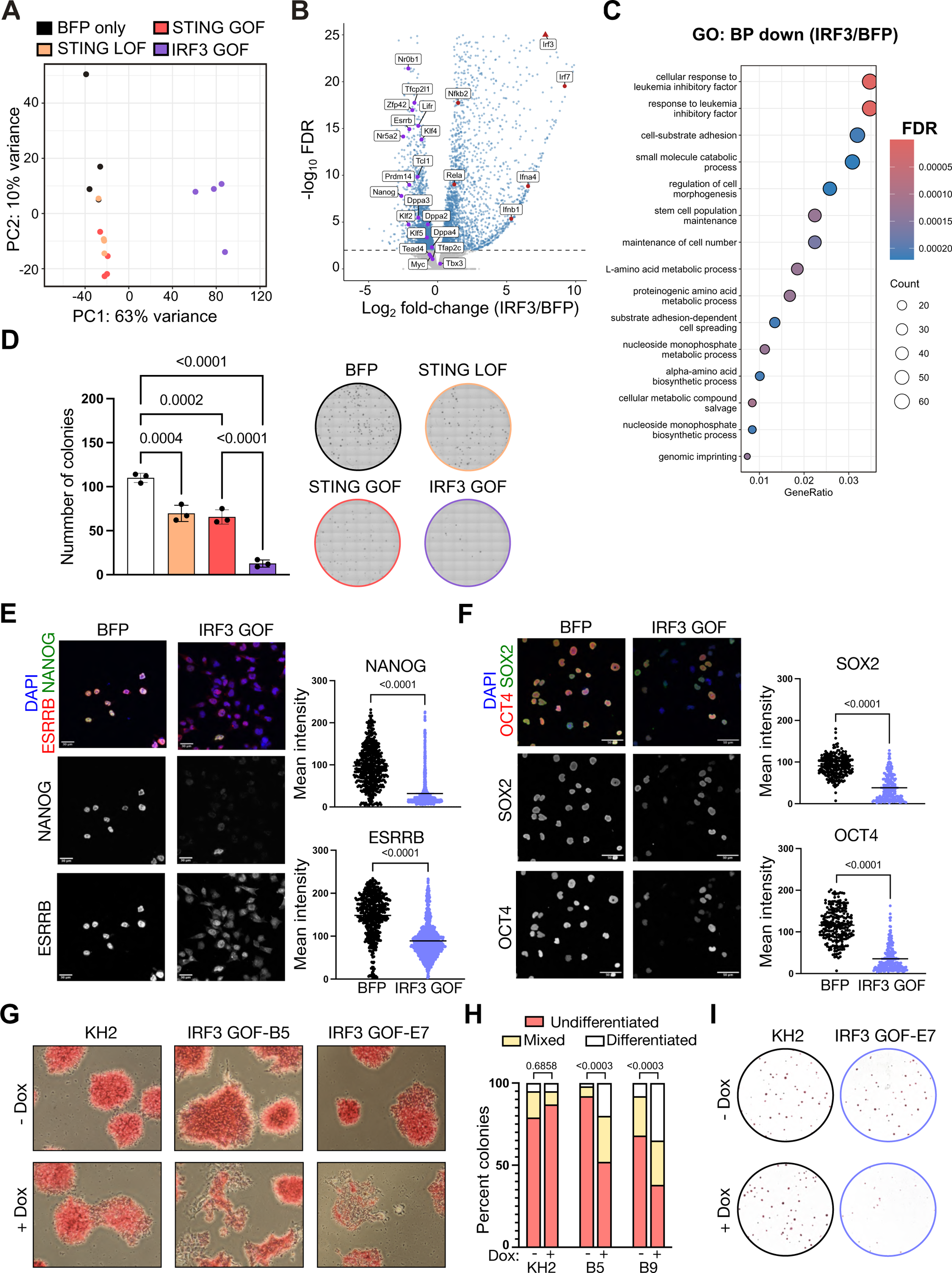
IRF3 activation is incompatible with pluripotency. a) PCA of all genes in the indicated ESC samples, showing that IRF3 GOF samples cluster separately from all other conditions along PC1. RNA-seq data are from 4 replicates from independent experiments. b) Volcano plot of differentially expressed genes between IRF3 GOF versus BFP-only transfected ESCs. Blue points are significantly differentially expressed between the conditions (FDR ≤ 0.01). A set of pluripotency genes are indicated in purple, and the majority are down-regulated in IRF3 GOF samples. Selected IFN-I genes are shown in red (triangle for IRF3 indicates −log_10_FDR >25). c) The top 15 biological processes (BP) enriched in GO analysis of significantly downregulated genes in IRF3 GOF over BFP-only conditions (FDR < 0.05). d) Colony formation assays in ESCs after transfection of the indicated constructs and sorting. ESCs were plated at low density and cultured in the presence of Dox for 5 days before fixing and staining for alkaline phosphatase. Data are mean +/- s.e.m for n=3 wells, representative of 3-4 experiments. P values, one-way ANOVA with Šídák’s multiple comparisons test. e) Examples and quantification of immunofluorescence experiments for ESRRB and NANOG, 36 h after transfection with BFP or IRF3 GOF and FACS. ESCs were plated onto Matrigel immediately after sorting and fixed 1h later. P values, Welch’s t-test. f) As in e, but staining for OCT4 and SOX2. g) Morphology and alkaline phosphatase staining of parental KH2 ESCs or stable IRF3 GOF ESCs (clone B5, E7) showing the latter begin to lose undifferentiated morphology by 72 h after Dox addition. Data are representative of 3 clones and 2 experiments. h) Quantification of ESC differentiation status in g), according to AP staining. Colonies were scored according to the criteria of undifferentiated: >80% AP-positive, mixed: 30-80% AP-positive, differentiated: <30% AP-positive. Data shown are from 2 experiments. P values, Chi-squared test with Bonferroni correction for multiple comparisons. i) Colony formation assay in KH2 parental ESCs or IRF3 GOF ESCs (clone E7) 5 days after plating at low density in the presence or absence of Dox. All data in figure are representative of at least 2 independent experiments unless otherwise indicated.

### The pluripotency network suppresses IFN-I signalling

The negative impact of IRF3 activation is unexpected given that ESCs are historically thought to be insensitive to interferon (Wang et al. 2014b; Wang et al. 2014c; Wu et al. 2018; Eggenberger et al. 2019). IFNβ, secreted upon IRF3/7 activation, binds to IFNAR receptors on nearby cells to activate JAK/STAT signalling. STAT1/2 bind to IFN-stimulated response elements (ISREs) together with IRF9, forming the ISGF3 complex. This complex induces transcription of IFN-I genes to amplify the innate immune response (**Fig.5A**). It has been shown that instead of canonical IFN-I signalling, ESCs constitutively express a set of ISGs that confer antiviral resistance, termed intrinsically expressed ISGs (IE-ISGs (Wu et al. 2018)). While IFNβ supplementation can increase ESC resistance to viruses, it induces only a subset of the ISGs induced in somatic cells (Wu et al. 2018; Muckenhuber et al. 2023). The reason for a dampened response in ESCs is still unknown. We next therefore investigated pathways downstream of IFNβ, to examine the wider consequences of IRF3 activation in ESCs. We first confirmed that JAK/STAT signalling is active in ESCs, with phospho-STAT1 induced in both ESCs and MEFs upon IFNβ addition (**Fig.5B**). However, ESCs treated with recombinant mouse IFNβ suffer no negative impact on ESC self-renewal (**Fig.5C**) – in contrast to the effects of IRF3 activation (**Fig.4D,I**) or to the antiproliferative effects of IFNβ seen in somatic cells (Hertzog et al. 1994). Culture of a large number of mouse embryos from morula to blastocyst stage in the presence of IFNβ also revealed no obvious abnormalities (**Fig.5D**), suggesting that blastocyst formation is not perturbed. In ESCs, we examined the expression of select canonical (*Il6, Tnf*) or IE-ISGs (*Isg15, Oas1a*) and confirmed that only the latter are significantly upregulated in ESCs (**Fig.5E-F**). Treating blastocysts with IFNβ suggested a similar pattern, with *Isg15* but not *Il6* upregulation detected (**Fig.S5A**). Thus, the negative impacts of IRF3 activation are clearly distinct to IFNβ treatment.

**Figure 5.**
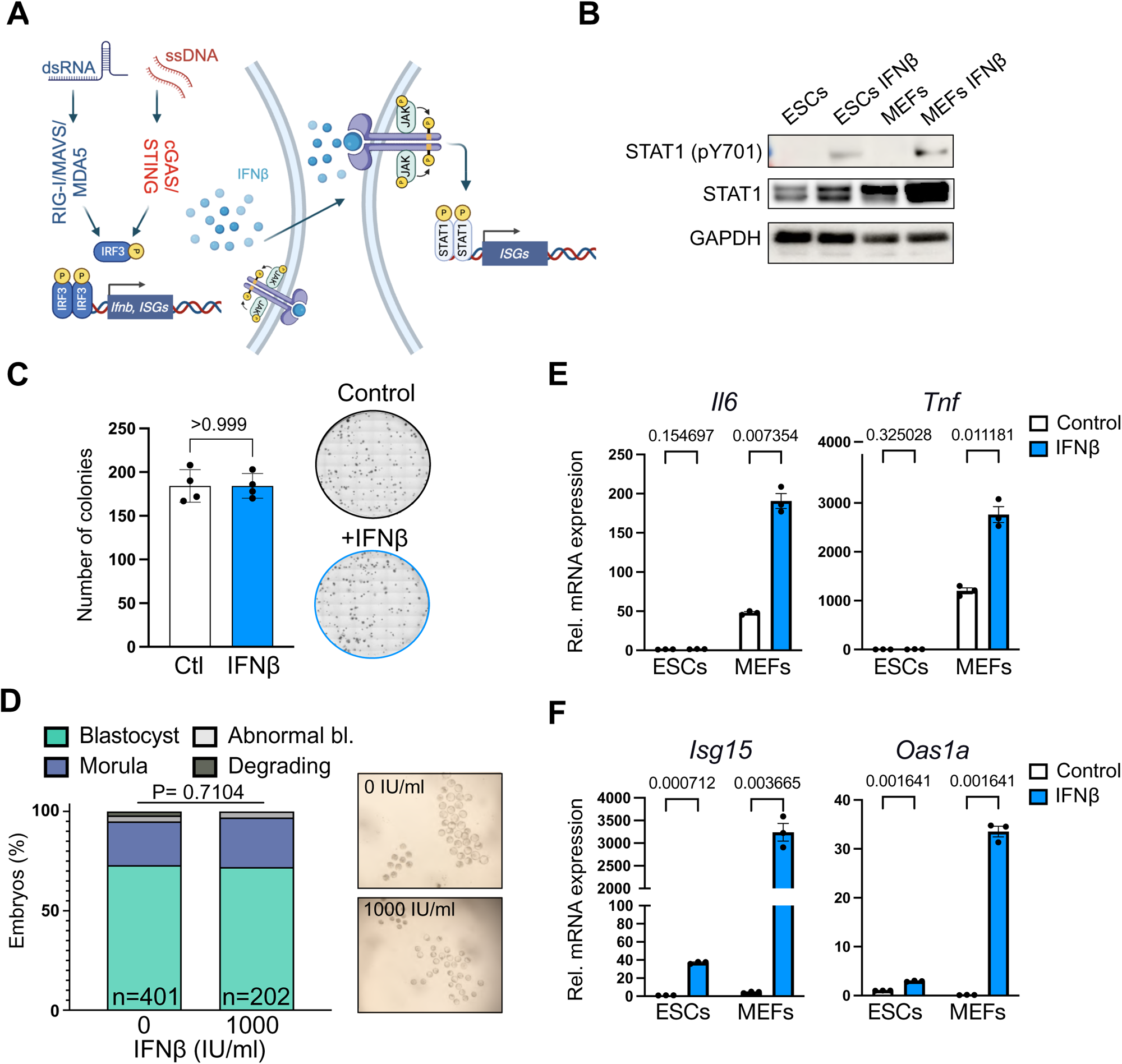
ESCs show a blunted response to Type I IFN. a) Simplified diagram of signalling pathways downstream of potential TE sensing and IRF3 activation. IRF3 activation leads to production of Type I IFNs (IFNβ, IFNα), which bind to IFNAR receptors on the plasma membrane of the same or neighbouring cells. IFN binding stimulates the JAK/STAT pathway, ultimately leading to STAT1/2 target gene expression, such as ISGs. Made with Biorender. b) Western blot showing STAT1 phosphorylation in both ESCs and MEFs 8 h after IFNβ treatment. c) Colony formation assay in ESCs in the presence or absence of exogenous IFNβ. Data are from 4 wells, representative of 3 experiments. P value, 2-tailed Student’s t-test. d) Example images and quantification of embryo progression (from morulae) after 24 h treatment with IFNβ. Data are combined from at least 3 experiments, n=embryo number. P value, Chi-squared test. e) RT-qPCR of canonical ISGs 8 h after IFNβ treatment in ESCs and MEFs. Data are mean +/- s.e.m, n=3 wells. P values, 2-tailed Student’s t-test with Holm-Šídák correction. f) As in e), but for intrinsically expressed ISGs (IE-ISGs), which show upregulation in ESCs. All data in the figure are representative of at least 2 independent experiments unless otherwise indicated.

To compare the global impact of IRF3 activation with IFNβ addition in ESCs and MEFs, we performed RNA-seq, integrating both datasets (**Fig.6A, 2D, S4B-C**). Significantly-upregulated genes upon IFNβ treatment in ESCs form only a small subset of MEF IFN targets, confirming a very limited IFN-I response (**Fig.6B, S5B**). GO analysis revealed that although interferon-related processes are upregulated in IFN-treated ESCs, no significant downregulated terms are found, in contrast to MEFs in which proliferation and cell cycle are impacted (**Fig.S5C**). Interestingly, our analysis revealed that IRF3 GOF substantially rescues downstream IFN-I gene activation in ESCs, inducing over a third of the genes activated in MEFs upon IFN (**Fig.6B**). These data indicate that IRF3 activation potentiates a much more widespread interferon response in ESCs than IFNβ treatment alone.

**Figure 6.**
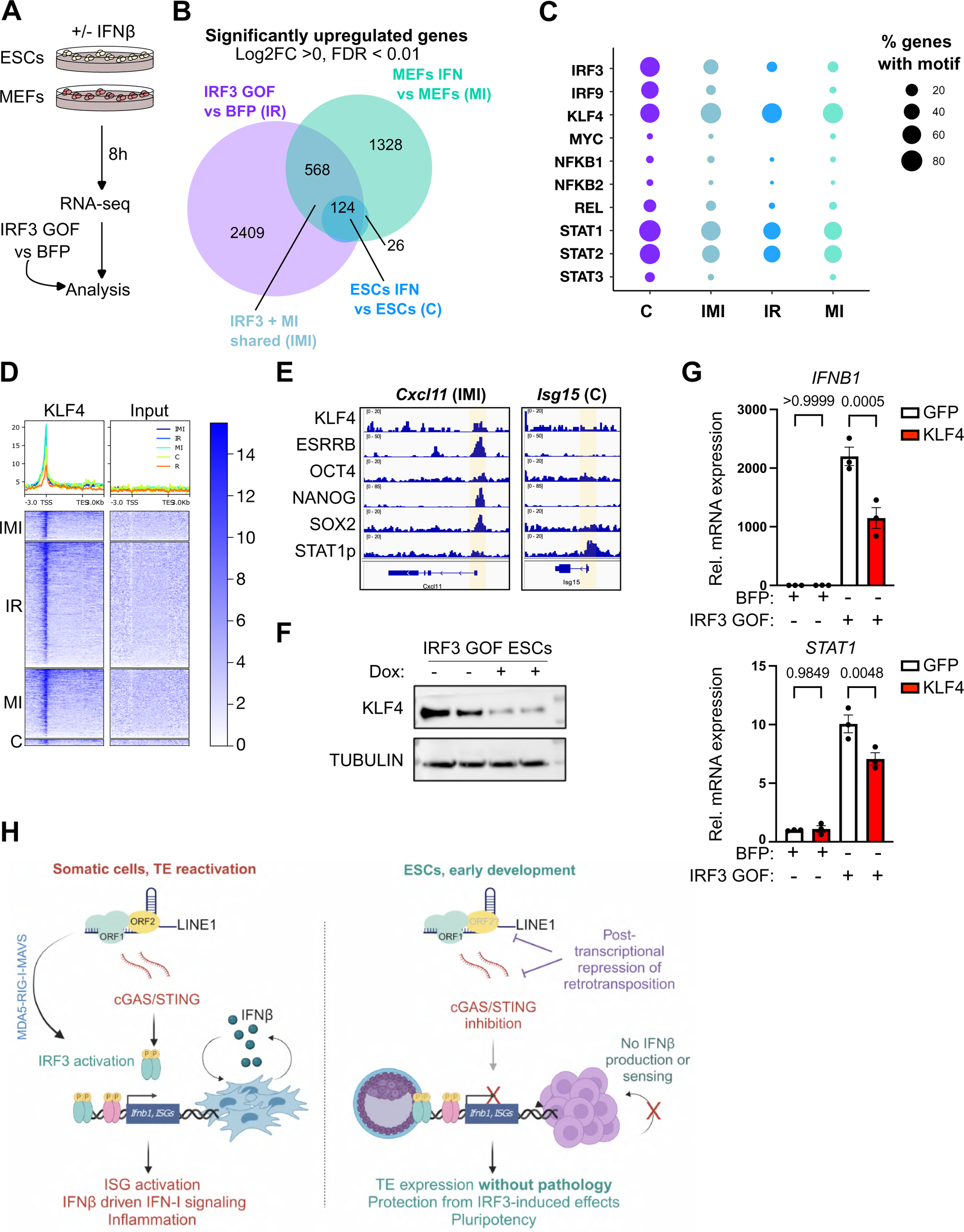
The pluripotency network antagonises IFN-I signaling networks. a) Diagram illustrating ESC/MEF IFNβ experiments for RNA-seq. Following sequencing, data were analysed together with IRF3 GOF data to compare direct IRF3 activation versus downstream IFN-I signalling. All samples for RNA-seq were collected from 4 independent experiments. b) Venn diagram showing the overlap in upregulated genes (FDR <0.01) between the indicated conditions in ESCs and MEFs. 2 genes were shared between IR and ESC+IFN (not shown). Gene sets are labelled with the corresponding acronym. c) Bubble plot displaying the percentage of genes in each gene set containing a significant motif occurrence for TFs of interest (q-value < 0.25). Motif scanning was performed +/- 1kb around the TSS of each gene and filtered for same-sense orientation. d) Spatial heatmaps displaying enrichment (RPGC) of KLF4 binding versus input (Aksoy et al. 2014) over gene bodies +/- 3kb. “R” refers to random gene set. The same result was seen for a second dataset (Fig.S6D)(Chronis et al. 2017). e) Example IGV browser screenshot showing transcription factor enrichments at the TSS (highlighted yellow) of selected genes from IMI and C gene sets. ChIP-seq datasets were reanalysed from (Chronis et al. 2017; Muckenhuber et al. 2023). f) Western blot of IRF GOF ESCs after 72 h with and without Dox-mediated induction, showing KLF4 downregulation. Data show samples from 2 experiments. g) RT-qPCR analysis of the indicated ISGs following co-transfection of pCW57.1 plasmids encoding BFP or BFP-t2a-IRF3 GOF with either pBABE-GFP or pBABE-KLF4 into 293T cells. Data are mean +/- s.e.m, n= 3 transfections. P values, one way ANOVA with Sidak’s correction for multiple comparisons. All data in the figure are representative of at least 2 experiments unless otherwise indicated. h) Model: in ESCs, but not somatic cells, high TE expression is uncoupled from pathology by two mechanisms. Suppression of cGAS/STING signalling and IRF3 phosphorylation prevents immune activation, while LINE1 retrotransposition is also post-transcriptionally suppressed. Rescuing a functional interferon response by IRF3 GOF antagonises pluripotency and self-renewal, highlighting the need for silencing TE sensing during development. Made with Biorender.

We next investigated how IRF3 rescues downstream IFN-I signalling. Cell-type specific activation at ISREs is well-known, and has been suggested to be partly due to chromatin state (Leviyang 2021; Muckenhuber et al. 2023). Analysis of IRF3/MEF+IFN shared genes (“IMI”) by ChromHMM (Ernst and Kellis 2017) revealed a decrease in accessibility and increased bivalency compared to upregulated genes common to all conditions (conserved, “C”) (**Fig.S6A, Table S2**). However, IMI genes also exhibit a higher basal expression level than C genes in ESCs (**Fig.S6B**). Thus, a more repressed chromatin state of IMI promoters likely does not fully explain their IFN insensitivity.

Given that IRF3 GOF causes widespread transcriptional activation at the same time as downregulating pluripotency, we next asked if such mechanisms might be linked. Indeed, NANOG has been suggested to directly inhibit NFKB activity (Torres and Watt 2008). Analysis of enriched motifs at the promoters of IMI, C, MEF only (MI) and IRF3 only (IR) genes (**Fig.6B**) revealed significant enrichment of STAT and IRF motifs as expected, but also of the pluripotency factor, KLF4 (**Fig.6C**). We examined published datasets for pluripotency factor binding at IMI genes and found evidence of KLF4, and to a lesser extent OCT4, SOX2 and cMYC, binding at these promoters in ESCs (**Fig.6D-E, S6C-D**). We reasoned that such occupancy might antagonise recruitment of ISGF3 (Stat1/2+IRF9); a repression that would be alleviated upon IRF3 activation and downregulation of pluripotency factors (**Fig.4E-F**), including KLF4 (**Fig.6F**). Indeed, KLF4 has been shown to interfere with cellular viral responses (Luo et al. 2016). In support of this, we found that KLF4 motifs are more centrally located over the TSS of gene sets not upregulated by IFNβ in ESCs (IR/IMI/MI). In contrast, the TSS of conserved (C) genes are more enriched for STAT1/2, IRF3 and IRF9 motifs (**Fig.S6E**). Notably, phospho-STAT1 binding is only robustly detected at C genes in ESCs upon IFNβ stimulation, supporting a potentially-weaker recruitment to other targets (**Fig.6E, S6C,F**). Testing our hypothesis, we overexpressed KLF4 at the same time as IRF3 GOF in a heterologous system and found that it can antagonise IRF3-mediated gene activation (**Fig.6G**). Our data suggest that weaker STAT1 binding, combined with KLF4 antagonism, prevents full ISG upregulation. Overall, these results point to a mutual incompatibility between pluripotency, cytosolic DNA sensing and Type I IFN signalling. Pluripotency serves to repress the activation of IRF3 and to limit Type I IFN responses, while activation of innate immunity has a strong, negative impact on early development.

## Discussion

The context of early development is uniquely associated with TE expression and yet apparently devoid of TE-induced pathology. Further, mammalian genomes provide ample historical evidence of retrotransposition in the germline and/or early embryo. Roughly 20% percent of cis regulatory elements are TE-derived (Sundaram et al. 2014), and insertions per individual have been estimated (Kano et al. 2009; Ewing and Kazazian 2010; Richardson et al. 2017). This raises the question: how is occasional TE activity enabled whilst still permitting TE expression – at the same time protecting the integrity of a developing embryo?

Here, we focused on two potential consequences of TE expression: sensing of TE reverse transcriptase activity, and TE retrotransposition itself. For the former, we reveal that the cGAS/STING pathway, the main mechanism linking sensing of cytosolic DNA to Type I IFN responses, is inhibited at multiple levels in ESCs. Rescue of this pathway by a constitutively active IRF3 is sufficient to trigger IFNβ production, downstream IFN-I responses, and destabilise pluripotency and self-renewal. Notably, this response is clearly distinct from the effects of direct IFNβ treatment. As reported previously (Wang et al. 2014b; Wang et al. 2014c), IFNβ only induces a very small subset of genes in ESCs, and we additionally show no effect on ESC self-renewal or blastocyst viability. We suggest that this is due – at least in part - to the pluripotency network suppressing IFN-driven gene activation.

An inhibitory link between the innate immune system and pluripotency has been previously suggested but never explained. For example, overexpression in human ESCs of a constitutively active IRF7 – a lymphoid-specific factor (Ma et al. 2023) - induces a mild differentiation defect, while IRF7-target genes were repressed in 293T cells upon co-expression of pluripotency factors (Luo et al. 2016; Eggenberger et al. 2019). This has limited relevance because IRF7 is not normally expressed in ESCs. In contrast, we show that IRF3 is ubiquitously expressed, including in ESCs, yet rendered inactive in development. Not only does IRF3 activation antagonise the pluripotency network, we suggest that pluripotency factors such as KLF4 in turn suppress ISGs. Such tightly-controlled, mutual inhibition serves to enforce suppression of innate immunity. It also provides mechanisms for IRF3-activated cells to be lost through differentiation or death in case of accidental triggering. It is important to note that in the context of the peri-implantation embryo, suppression of IFNβ production is even more important given the nearby maternal endometrium. The maternal endometrium is IFN responsive, with implantation itself known to be an inflammatory process (reviewed in (Yockey and Iwasaki 2018)). It is possible that activation of cGAS/STING and downstream IRF3 in the embryo would thus interfere with implantation and early post-implantation development. While this specific question has not been investigated, IFN-I dysregulation in the mother or mid-gestation foetus is associated with cases of developmental defects or pregnancy complications (Yockey and Iwasaki 2018).

This work builds upon and rationalises previous findings of development-specific strategies to deal with viral pathogens. This includes intrinsically-expressed ISGs that provide antiviral protection in the absence of PAMPs (Wu et al. 2018), as well as antiviral RNAi (Flemr et al. 2013; Maillard et al. 2013; Maillard et al. 2016; Poirier et al. 2021). At the same time, MAVS downregulation in ESCs contributes to their insensitivity to dsRNA (Witteveldt et al. 2019). The existence of alternative defence mechanisms is consistent with an absolute requirement to prevent canonical IFN-I activation in development. We argue that this requirement may be due to the need to avoid responding to TEs. Although we cannot rule out other reasons such as avoiding a response to a sporadic viral infection, the high expression of many TEs in development poses a significant endogenous threat even in the absence of pathogens. Indeed, SINE elements are abundant in ESCs, and can form dsRNA (Babiarz et al. 2008). An elevated presence of dsRNA has been recently reported in ESCs, including an enrichment for TE-containing genes (Witteveldt et al. 2025). The reverse transcriptase activities of both LINE1 and ERVs are moreover capable of generating ssDNA and inducing inflammation (Kassiotis and Stoye 2016; Thomas et al. 2017; De Cecco et al. 2019). Although rare, germline or early embryo retrotransposition is a core driver of mammalian genome complexity (Sundaram and Wang 2018). A canonical IFN response only emerges sometime later in development or differentiation (D’Angelo et al. 2016), by which time TEs are also downregulated. The exact timings of IFN-I rescue remain to be discovered, and it will be interesting in future work to understand when in development cGAS/STING signalling is reactivated. Moreover, future studies should uncover the exact mechanisms through which cGAS/STING expression, or IRF3 phosphorylation, are inhibited in ESCs. Both cGAS/STING itself as well as the TBK1-IRF3 relationship are frequently disrupted in cancer or upon infection (Du et al. 2015; Yu et al. 2022; Sasaki et al. 2023). Any findings in development may thus also illuminate relevant disease contexts. Finally, our work raises the question of the existence of similar silencing mechanisms in other stem cell or progenitor populations, including those with reported TE expression (Chapman et al. 1984; Muotri et al. 2005; Coufal et al. 2009; Golding et al. 2010).

As well as investigating TE sensing, we examined control of TE activity in ESCs, focusing on LINE1. We show that despite very high levels of LINE1 expression – much greater than in senescent cells – cytosolic DNA is not detected in ESCs, and LINE1 retrotransposition is significantly repressed. This is supported by IP-MS of ORF1p in ESCs, which pulls out many known interactors of LINE1 found in other cell types yet no enrichment of proteins that bind LINE1 retrotransposition or ssDNA intermediates. Our results highlight a clear uncoupling of LINE1 expression and activity in early development, and point to pluripotency-specific, post-transcriptional mechanisms to inhibit retrotransposition. As with tRNA fragments that have been reported to inhibit IAP and MusD LTR reverse transcriptase (Schorn et al. 2017), similar mechanisms may restrict LINE1 activity in ESCs. Although DICER and miRNA activity has been suggested to inhibit retrotransposition in pluripotent cells (Hamdorf et al. 2015; Bodak et al. 2017; Tristan-Ramos et al. 2020), we found effects of DICER ablation upon LINE1 retrotransposition in mouse ESCs to be fairly mild (data not shown). It will be critical in future studies to discern how LINE1 activity is uniquely repressed in development, for example at the level of regulating ORF2p and/or target-primed reverse transcription. Such pathways may be furthermore targetable in disorders where TE activity needs to be prevented. During development, we suggest that the combination of suppressed TE sensing and suppressed TE activity (**Fig.6H**) provides an environment where TE expression can be safely uncoupled from the harmful impacts of retroelement reactivation.

## Materials and Methods

### Embryo culture

All animal experiments were performed according to a UK Home Office Project License in a Home Office-designated facility using 4- to 6-week-old female and 2- to 6-months-old male C57Bl/6J mice purchased from Envigo. Animals were maintained on a 12-h light/dark cycle and provided with ad libitum food and water in individually ventilated cages. Female mice were superovulated by intraperitoneal injection of 5 IU of pregnant mare serum gonadotropin (PMSG; Folligon, MSD Animal Health), followed by 5 IU of human chorionic gonadotropin (hCG; Chorulon, MSD Animal Health) 46–48h later, and then placed immediately with males. Presence of a copulatory plug the following morning was taken to indicate mating had occurred, and the day of plug was designated embryonic day (E) 0.5. Embryos were recovered from the uterus at morula (E2.5) or blastocyst (E3.5) stage by flushing in M2 medium (Merck), and cultured in pre-equilibrated KSOMaa medium (Merck) in a humidified incubator at 37 degrees and 5% CO_2_. For some experiments, morulae were stored in LN_2_ after slow process cryopreservation in straws containing a M2/Propan-2-ol solution and a M2/sucrose solution. Embryos were thawed by removal from LN_2_ to room temperature for 1 minute, then straw contents were released to M2/sucrose for 5 minutes, M2 for 7 minutes, and finally KSOMaa for ongoing culture. Embryos were cultured in 1000 IU/ml recombinant mouse IFNβ (Biotechne) for 24 hours from E2.5 to E3.5, or for 6 hours from approximately 10am to 4pm on E3. For RNA collection, 15-26 embryos per replicate were lysed in 75 µL RLT buffer before RNA extraction with the RNeasy Mini kit (Qiagen) as below.

### Cell culture

Mouse E14Tg2A (E14) ES cells (male) were cultured on 0.1% gelatine-coated plates in serum/LIF culture medium (high glucose DMEM GlutaMAX (Thermo Fisher Scientific, USA), 10% heat inactivated FBS (Thermofisher, USA) 0.1 mM Beta-mercaptoethanol (Millipore) and 1,000U/ml of LIF supplement (ESGRO, Millipore). Medium was changed every day and cells passaged every other day using TrypLE (Gibco). For spontaneous differentiation experiments, 200,000 cells were plated in non-LIF supplemented media for 5 days in 10cm plates, changing media every day. Mouse KH2 ES cells (Beard et al. 2006) were cultured on 0.1% gelatine-coated plates in serum/LIF culture medium, supplemented with 1 μM of PD0325901 and 3 μM of CHIR99021.

MEF cells and HEK 293T cells were cultured in 10% serum DMEM media (Thermo Fisher Scientific). For senescence induction, MEFs were grown until 70% confluence and treated with 20 μM etoposide for 3 days, washed with PBS and cultured in MEF media for 60-72 days supplemented with Anti-anti (Gibco), renewing the media every 3 days. Senescence was confirmed with a senescence beta-galactosidase staining kit (Cell Signaling), according to the manufacturer’s protocol. All cells in this study were routinely tested and found to be mycoplasma negative using a MycoAlert biochemical luminescence kit (Lonza).

### STING/IRF3 constructs

STING GOF mutations (V155M in human, V154M in mouse (Bouis et al. 2019)) have been described to cause an autoinflammatory disease named STING-associated vasculopathy with onset in infancy (SAVI)(Cerboni et al. 2017). The STING HAQ loss of function allele contains 3 non-synonymous single nucleotide substitutions: R71H, G230A and R293Q in human (C71H, I229A, R292Q in mouse)(Patel et al. 2017), which we incorporated on top of a V154M background in the STING LOF mutant. IR3 GOF constructs were generated by replacing 5 Serine/Threonine residues from the C-terminal IRF3 transactivation domain by the phosphomimetic, aspartic acid, which makes IRF3-5D (Lin et al. 1999; Wang et al. 2014a). For transient STING/IRF3 overexpression experiments, STING GOF, STING LOF, and IRF3 GOF constructs were designed to be expressed downstream of BFP and a t2a cleavage element and synthesised (Genscript) then subcloned into the Dox-inducible expression vector, pCW57.1, (a gift from David Root; Addgene plasmid #41393, http://n2t.net/addgene:41393, RRID:Addgene_41393).

### Transfections

Transfections were performed into cells using 3μl/µg DNA of Lipofectamine 2000 according to manufacturer’s procedures. The medium was changed next day, including addition of Dox where indicated. For transfection of 2’3’cGAMP and G3-YSD/YSD-C ligands (Invivogen), reagents were reconstituted in water at a stock concentration of 1 mg/ml. 8 µg of 2’3’ cGAMP and 1 µg of G3-YSD ligands with the corresponding controls were transfected per well in a 12-well plate using Lipofectamine 2000, using the same ratios as for DNA transfections (Thermo Fisher Scientific, USA). For STING/IRF3 transfections, 8 μg of plasmid was transfected in suspension into 1 million cells in a 10 cm dish with 1μg/ml of doxycycline, which was renewed with every media change. 36h hours later, BFP-positive cells were sorted and processed for downstream analyses (RT-qPCR, RNA-seq, immunofluorescence). For KLF4/IRF3 overexpression, pBABE-based expression vectors for KLF4 (a gift from Daniel Peeper (Rowland et al. 2005)), or GFP were transiently transfected into 293T cells together with pCW57.1-IRF3-GOF or pCW57.1-BFP. 24 h later, 1 μg/mL Dox was added and samples harvested for RNA analysis after a further 24 h.

For stable STING/IRF3 overexpression, STING constructs were synthesised without BFP, for subcloning into pCol-TGM, for targeting into the *Col1a1* locus of KH2 ESCs (Beard et al. 2006). IRF3 GOF constructs were synthesised similarly, with or without a downstream t2a-mCherry to monitor expression. pCol-TGM was a gift from Scott Lowe (Addgene plasmid # 32715). KH2 ESCs were co-transected with the relevant pCol-TGM construct together with a Flippase plasmid, then later selected with 150 μg/mL hygromycin for 7 days before picking colonies, genotyping, and expanding. Multiple clones per construct were tested and validated for experiments. Dox-induction was performed at 1-2 μg/mL dose for the indicated time points, refreshing Dox every 24 h.

### Interferon treatment

ESCs, MEFs or embryos were treated with 200 or 1000 IU of mouse IFNβ at different timepoints (Biotechne). 100 µg/ml IFNβ storage stocks (equivalent to 1.2*10^8^IU/ml) were prepared in 0.1% PBS BSA and used to make a 1000X working stock in 0.1% PBS BSA. The 1X working solution was added fresh to culture medium.

### LINE1 reporter assays

800,000 ESCs or 293T cells were plated on 10 cm plates the day before and transfected with 4.5 µg of LINE1 GFP reporter plasmid (L1 ORFeus, (An et al. 2006)) and 0.5 µg of mCherry-expressing plasmid with Lipofectamine 2000 reagent (Thermo Fisher Scientific, USA). Media was changed just before transfection, as well as 24 and 48 hours post transfection. Individual plasmids were transfected separately for compensation controls. Samples were later collected and resuspended in FACS buffer (DMEM - phenol-red free, 3% FBS, 3 mM EDTA, 1:8000 sytox blue) and assessed by flow cytometry, 72 hours post transfection. Data were collected on a BD LSR II and analysed using FlowJo v10.10.0.

### Colony formation assays

500 or 1000 ESCs per well were plated per well of a gelatin-coated 12 or 6-well-well plate, respectively. ESCs were kept in culture 5 days with daily renewals of media +/- 1 µg/ml Dox or IFNβ where relevant. On the last day, ESCs were fixed with 4% PFA and stained with an Alkaline phosphatase kit (Vector, USA) according to the manufacturer’s protocol.

### Immunofluorescence

ESCs and MEFs were plated on matrigel-coated 8-chamber coverslips (Thermofisher, USA) for 2 h and fixed for 15’ with 4% paraformaldehyde (Thermofisher, USA) in Phosphate Buffered Saline (PBS) at room temperature (RT). To permeabilize, different concentrations of Triton were used depending on the staining, with the standard condition being 0.1% triton for 10’, (Table S3). Where relevant, ESCs were first sorted by FACS according to BFP fluorescence before pelleting, resuspending in ESC media and seeing on chamber slides as above. Samples were then blocked for 30’ in blocking buffer, then incubated overnight in a humidified compartment at 4°C to prevent the samples from drying, with primary antibody (Ab) in blocking solution at the specified dilutions (Table S3). Samples were washed 3 times the next day with PBS, incubated with the appropriate secondary antibodies for 45’ at RT protecting samples from light, washed 3 times with PBS, then mounted in Vectashield solution (Thermofisher, USA) with 4′,6-diamidino-2-phenylindole (DAPI) as counterstaining. Stained samples were imaged using a confocal laser-scanning microscope (Leica TCS SP5) with 40x dry and 63x oil immersion objective lenses and processed using FIJI. Quantifications were performed with CellProfiler (ORF1p quantification, colony formation assays) or Qpath (all other experiments).

Staining of ssDNA was performed as described by (De Cecco et al. 2019). Cells were seeded on Matrigel coated coverslips and fixed on ice with 4% PFA for 10’ and then incubated in 100% methanol at −20°C overnight. The cells were then incubated with 200 μg/ml RNase A at 37°C for 4h in gentle agitation. Cells were later blocked in 3% BSA solution for 1h and incubated overnight at 4°C with the relevant mouse anti ssDNA (Enzo, F7-26) primary antibody diluted in 3% BSA.

### Western blot

Whole cell extracts were prepared in ice-cold RIPA buffer (50 mM Tris pH8, 1 mM EDTA, 0.5 mM EGTA, 1% Triton X10, 0.1% sodium deoxycholate, 140 mM NaCl) containing Halt protease inhibitors (Thermofisher, USA). Extracts were incubated in a rotating wheel at 4°C for 15’, followed by a 14,000rpm 15’ centrifugation step to pellet debris. Samples were quantified using the Pierce BCA protein assay (Thermofisher, USA) and 20-40μg of total protein was loaded per sample. Proteins were separated in a Bolt Bis-Tris Plus 4-12% gel, using Prestained protein plus ladder (Thermofisher, USA). After this step, a dry transfer was carried out onto a PVDF membrane using the iBlot2 Dry blotting system (Thermofisher, USA). The membrane was then blocked for an hour in 5% w/v powder milk in TBST. Membranes were then incubated overnight at 4°C in primary antibody (Table S3). The next day membranes were washed in Tris-buffered saline TWEEN 0.2% (TBST) and incubated in the appropriate anti-rabbit/goat secondary antibodies conjugated to HRP. Membranes were imaged in an Amersham imager 680 (Cytivia, UK). Quantification was performed using the software ImageJ.

### Co-immunoprecipitations and IP-mass spectrometry (IP-MS)

Whole cell extracts were prepared from ESCs by scraping plates with ice-cold RIPA buffer containing protease inhibitors, prepared as for WB and quantified using Pierce BCA protein assay kit. Protein A magnetic beads were prepared with 2.5 μg the appropriate antibodies: ORF1(Abcam, #ab216324), anti-MOV10 (Abcam, #ab80613) and control Rabbit IgG (Cell Signalling Technology, #2729) as negative control. 30 μl of protein A Dynabeads were bound to antibodies by incubating them with 2.5 μg of Ab in 200 μl of RIPA buffer for 20 minutes at room temperature. The beads were then crosslinked to the antibody using 5 mM BS3 in conjugation buffer (20 mM NaH_2_PO_4_, 0.15 M NaCl in H_2_O) for 30 minutes at room temperature. Beads bound to the appropriate antibodies were used to immunoprecipitate the proteins of interest. 1 mg of total protein extract plus inhibitors was incubated overnight in a rotating wheel. Next day immunocomplexes bound to the beads were washed first in RIPA and then twice for 1 minute in TSE buffer (1% triton 2mM EDTA, 20mM EDTA, 20mM Tris pH8 and150mM NaCl). Samples were eluted by boiling 5’ at 90°C in LDS buffer + 2mM DTT or elution buffer (mass-spectrometry: 50mM HEPES ph8, 1% (wt/vol) SDS, 1% (vol/vol) Triton X-100, 1% (vol/vol) NP-40, 1% (vol/vol) Tween 20, 1% (wt/vol) deoxycholate, 5 mM EDTA, 50 mM NaCl, 1% (vol/vol) glycerol. For IP-MS, samples were processed with SP3 and digested with trypsin. Each sample was run in 2 technical replicates on the spectrometer (HFX). Data were processed using the MaxQuant software, statistical analysis and visualizations performed in Perseus. A non-parametrical Welch t-test was used for the statistical analysis.

### RT-qPCR

Total RNA was extracted from cell pellets using the RNeasy Mini kit (Qiagen) according to manufacturer’s procedures, after washing with PBS then lysis in RLT buffer plus 2-mercaptoethanol. Up to 1 μg of RNA in a maximum volume of 9 μl was used for cDNA generation using the High-capacity RNA to cDNA kit (Thermo Fisher, USA) according to the manufacturer’s protocol. For experiments involving sorted cells and embryos, the RNeasy Micro kit was used according to manufacturer’s procedures. In this case the whole eluate was used for cDNA generation. qPCR was performed using SYBR green kit (KAPA, Roche) on a QuantStudio5 (Thermo Fisher Scientific, USA) in 384 well plates (Thermo Fisher scientific, USA), using up to 4 ng of cDNA per well. Each reaction per primer pair (Table S3) was run in triplicates per each sample. The arithmetic mean of *Rpl7* and *H2A* housekeeping genes was used to normalize the samples. Primers were used at 200 nM final concentration.

### RNA-seq

RNA from sorted BFP+ cells transfected with the relevant construct and treated with Dox for 36h was extracted using the Qiagen RNA micro kit. RNA from MEF and ESC samples for interferon experiments was extracted using the RNeasy Mini Kit. RNA quality control was confirmed using a bioanalyzer before sending for sequencing (Novogene, PE 150bp, approx. 25M reads per sample. RNA-sequencing data are provided in **Table S1**.

### RNA-seq data processing and analysis

RNA-seq data were processed using the nf-core/rnaseq pipeline (Ewels et al. 2020; Patel et al. 2024b) (nfcore/rnaseq v3.14.0; Nextflow v24.10.3). Raw RNA sequencing reads (PE150) were trimmed using TrimGalore (Krueger 2021) (v0.6.7) with default settings. Data were aligned to the mm39 mouse reference genome using STAR (Dobin et al. 2013) (v2.7.9a; ‘--twopassMode Basic’) and gene-level quantification was performed using Salmon (Patro et al. 2017) (v1.10.1; default options) relative to Ensembl annotation (v110). Principle component analysis (PCA) was performed on count matrices normalized by variance stabilizing transformation (VST). Differential expression analysis was performed using DESeq2 (Love et al. 2014) (v1.44.0) to identify DEGs (FDR < 0.01) from the following comparisons: 1) IRF3 GOF ESCs and BFP-only control ESCs, 2) STING GOF ESCs and BFP-only control ESCs, 3) STING LOF ESCs and BFP-only control ESCs, 4) IFNβ-exposed ESCs vs untreated control ESCs, 5) IFNβ-exposed MEFs vs untreated control MEFs. Variance partitioning suggested the presence of batch effects between ESC BFP-only control replicates, which were corrected through the RUV-s method (Risso et al. 2014) (RUVSeq v) and incorporated into the DESeq design of relevant comparisons. Gene ontology analysis was performed using clusterProfiler (Wu et al. 2021) (v4.12.6) to identify significantly enriched biological processes in DEGs from each comparison (q-value < 0.05). To measure and plot the GSEA enrichment profile presented in Figure S4A, results from differential expression analysis between IRF3 GOF ESCs and BFP-only control ESCs were ranked (log2 fold-change) and used as input for GSEA analysis run with clusterProfiler and plotted with enrichplot (v1.24.2) (Yu 2025).

### Integration and analysis of published datasets

To establish chromatin regulatory landscapes and relevant transcription factor binding patterns, we downloaded and analysed published ChIP-seq and Atac-seq data in mouse ESCs and MEFs from the ENCODE Portal (Consortium 2012; Luo et al. 2020) and GEO (Barrett et al. 2013) (**Table S2,** (Aksoy et al. 2014; Yue et al. 2014; Chronis et al. 2017; Sethi et al. 2020; Muckenhuber et al. 2023)). ChIP-seq and Atac-seq data were processed using the nf-core/chipseq (v2.1.0-g76e2382) and nf-core/atacseq (v2.1.2) pipelines, respectively (Ewels et al. 2020; Patel et al. 2023; Patel et al. 2024a). Raw fastq were trimmed using TrimGalore (default settings). Mapping to the mm39 genome was performed with BWA (Li and Durbin 2009) for ChIP-seq and Atac-seq (v0.7.18-r1243-dirty; default settings). In ChIP-seq, duplicates were marked and removed (Picard v3.2.0-1-g3948afb6b) from input controls but were maintained in target datasets. Reads with MAPQ below 10 were discarded and reads annotated to mm39 blacklisted regions (Ogata et al. 2023) were removed. Normalized bigwig files were generated using deepTools bamCoverage (Ramírez et al. 2016) (v3.5.3; ‘--normalizeUsing RPGC’). Tracks were visualized in the IGV genome browser (Robinson et al. 2011). To produce the heatmaps and profiles in Figures 6D, S6D, and S6F, normalized signal was extracted over specified gene regions using deepTools computeMatrix (‘reference-point --beforeRegionStartLength 3000 -- regionBodyLength 5000 --afterRegionStartLength 3000’) followed by data visualization using plotHeatmap or plotProfile.

### Chromatin state characterization and fold-enrichment analysis

We ran ChromHMM (Ernst and Kellis 2012) (v1.26) on ChIP-seq and Atac-seq data in mouse ESCs for the following targets: H3K4me3, H3K4me1, H3K27ac, H3K9me3, H3K36me3, H3K27me3, and Atac-seq (**Table S2** (chromHMM)). Analysis was performed at the default 200-bp resolution using BAM files and relevant control files aligned to the mm39 genome (BinarizeBam; default parameters). Models were generated at multiple state-level resolutions ranging between 12 and 22-states (LearnModel; default parameters). The 14-state model was chosen as it presented interpretable epigenetic states with distinct mark combinations. Chromatin states were manually annotated in relation to evaluated metrics, as described in Figure S6A and **Table S2**. The ChromHMM OverlapEnrichment function was used to calculate fold-enrichment scores for chromatin states over genomic features and over regions centered on C/IR/IMI/MI promoters (2kb +/- TSS). Enrichment values over C/IR/IMI/MI gene sets were log2-transformed and are displayed in Figure S6A (left).

### Motif analyses

Motifs enriched in promoter regions of C/IR/IMI/MI gene sets were identified using the AME tool (McLeay and Bailey 2010) of the MEME Suite (Bailey et al. 2015) (v5.5.7) and the HOCOMOCO Mouse (v11 CORE) database (Vorontsov et al. 2024). The promoter region was chosen for motif detection due to the expected elevated enrichment at TSS of our selected transcription factors, which we observed in our ChIP-seq analysis. Following AME, relevant motifs were selected based upon motif enrichment clustering patterns, relative enrichment in the gene sets, and expression of the transcription factors. The FIMO tool of the MEME Suite (Grant et al. 2011) was then used to extract locations of the chosen motifs in C/IR/IMI/MI gene sets. To compare motif prevalence, FIMO was performed using promoter regions (+/- TSS 1kb) and the percentage of genes in each set containing a significant hit for each motif were calculated (**Fig.6C**). To identify the relative locations of transcription factor binding sites, FIMO was performed using a broader region (+/- TSS 3kb) of each gene set. Locations of significant hits were standardized and strand-normalized relative to gene features, and density curves were plotted to display motif distributions (**Fig.S6E**).

### Statistical Analysis

all experiments were performed multiple times with the equivalent results. Data in individual graphs are shown either combined from multiple experiments, or from biological replicates from one experiment, representative of at least 1-2 more experiments. Details of replication and all statistical tests are provided in the relevant figure legends, including which corrections were performed for multiple testing or unequal variance.

## Acknowledgements

We thank Aydan Bulut-Karslioglu, Matthias Merkenschlager, Marco Trizzino, Juanma Vaquerizas, and members of the Percharde lab for project input and/or critical reading of the paper. This work was supported by a UKRI Future Leaders Fellowship (MC_EX_MR/X022560/1) and MRC core funding (MRC; MC_UP_1605/4) to M. Percharde.

## Author Contributions

Conceptualization: MP

Methodology: MP, FG-L, AP, BJL

Investigation: FG-L, AP, AC, BJL, KS, ARG, BM

Visualization: MP, AP, FG-L

Supervision: MP, ARB, JG

Writing—original draft: MP, FG-L

Writing—review & editing: All authors

## References

Aksoy I, Giudice V, Delahaye E, Wianny F, Aubry M, Mure M, Chen J, Jauch R, Bogu GK, Nolden T et al. 2014. Klf4 and Klf5 differentially inhibit mesoderm and endoderm differentiation in embryonic stem cells. Nat Commun 5: 3719.

Almeida MV, Vernaz G, Putman ALK, Miska EA. 2022. Taming transposable elements in vertebrates: from epigenetic silencing to domestication. Trends Genet 38: 529–553.

An W, Han JS, Wheelan SJ, Davis ES, Coombes CE, Ye P, Triplett C, Boeke JD. 2006. Active retrotransposition by a synthetic L1 element in mice. Proc Natl Acad Sci U S A 103: 18662–18667.

Andzinski L, Wu CF, Lienenklaus S, Kroger A, Weiss S, Jablonska J. 2015. Delayed apoptosis of tumor associated neutrophils in the absence of endogenous IFN-beta. Int J Cancer 136: 572–583.

Ardeljan D, Steranka JP, Liu C, Li Z, Taylor MS, Payer LM, Gorbounov M, Sarnecki JS, Deshpande V, Hruban RH et al. 2020a. Cell fitness screens reveal a conflict between LINE-1 retrotransposition and DNA replication. Nat Struct Mol Biol 27: 168–178.

Ardeljan D, Wang X, Oghbaie M, Taylor MS, Husband D, Deshpande V, Steranka JP, Gorbounov M, Yang WR, Sie B et al. 2020b. LINE-1 ORF2p expression is nearly imperceptible in human cancers. Mob DNA 11: 1.

Babiarz JE, Ruby JG, Wang Y, Bartel DP, Blelloch R. 2008. Mouse ES cells express endogenous shRNAs, siRNAs, and other Microprocessor-independent, Dicer-dependent small RNAs. Genes & development 22: 2773–2785.

Bailey TL, Johnson J, Grant CE, Noble WS. 2015. The MEME Suite. Nucleic Acids Research 43: W39–W49.

Barrett T, Wilhite SE, Ledoux P, Evangelista C, Kim IF, Tomashevsky M, Marshall KA, Phillippy KH, Sherman PM, Holko M et al. 2013. NCBI GEO: archive for functional genomics data sets—update. Nucleic Acids Research 41: D991–D995.

Beard C, Hochedlinger K, Plath K, Wutz A, Jaenisch R. 2006. Efficient method to generate single-copy transgenic mice by site-specific integration in embryonic stem cells. Genesis 44: 23–28.

Bestor TH, Bourc’his D. 2004. Transposon silencing and imprint establishment in mammalian germ cells. Cold Spring Harb Symp Quant Biol 69: 381–387.

Bodak M, Cirera-Salinas D, Yu J, Ngondo RP, Ciaudo C. 2017. Dicer, a new regulator of pluripotency exit and LINE-1 elements in mouse embryonic stem cells. FEBS Open Bio 7: 204–220.

Bouis D, Kirstetter P, Arbogast F, Lamon D, Delgado V, Jung S, Ebel C, Jacobs H, Knapp AM, Jeremiah N et al. 2019. Severe combined immunodeficiency in stimulator of interferon genes (STING) V154M/wild-type mice. J Allergy Clin Immunol 143: 712–725 e715.

Brouha B, Schustak J, Badge RM, Lutz-Prigge S, Farley AH, Moran JV, Kazazian HH, Jr. 2003. Hot L1s account for the bulk of retrotransposition in the human population. Proc Natl Acad Sci U S A 100: 5280–5285.

Burns KH. 2017. Transposable elements in cancer. Nat Rev Cancer 17: 415–424.

Carpenter S, O’Neill LAJ. 2024. From periphery to center stage: 50 years of advancements in innate immunity. Cell 187: 4429–4430.

Castaneda J, Genzor P, Bortvin A. 2011. piRNAs, transposon silencing, and germline genome integrity. Mutat Res 714: 95–104.

Cerboni S, Jeremiah N, Gentili M, Gehrmann U, Conrad C, Stolzenberg MC, Picard C, Neven B, Fischer A, Amigorena S et al. 2017. Intrinsic antiproliferative activity of the innate sensor STING in T lymphocytes. J Exp Med 214: 1769–1785.

Chapman V, Forrester L, Sanford J, Hastie N, Rossant J. 1984. Cell lineage-specific undermethylation of mouse repetitive DNA. Nature 307: 284–286.

Chen JM, Stenson PD, Cooper DN, Ferec C. 2005. A systematic analysis of LINE-1 endonuclease-dependent retrotranspositional events causing human genetic disease. Hum Genet 117: 411–427.

Chen LL, Yang L, Carmichael GG. 2010. Molecular basis for an attenuated cytoplasmic dsRNA response in human embryonic stem cells. Cell Cycle 9: 3552–3564.

Chiappinelli KB, Strissel PL, Desrichard A, Li H, Henke C, Akman B, Hein A, Rote NS, Cope LM, Snyder A et al. 2015. Inhibiting DNA Methylation Causes an Interferon Response in Cancer via dsRNA Including Endogenous Retroviruses. Cell 162: 974–986.

Chronis C, Fiziev P, Papp B, Butz S, Bonora G, Sabri S, Ernst J, Plath K. 2017. Cooperative Binding of Transcription Factors Orchestrates Reprogramming. Cell 168: 442–459 e420.

Consortium EP. 2012. An integrated encyclopedia of DNA elements in the human genome. Nature 489: 57–74.

Coufal NG, Garcia-Perez JL, Peng GE, Yeo GW, Mu Y, Lovci MT, Morell M, O’Shea KS, Moran JV, Gage FH. 2009. L1 retrotransposition in human neural progenitor cells. Nature 460: 1127–1131.

D’Angelo W, Acharya D, Wang R, Wang J, Gurung C, Chen B, Bai F, Guo YL. 2016. Development of Antiviral Innate Immunity During In Vitro Differentiation of Mouse Embryonic Stem Cells. Stem Cells Dev 25: 648–659.

De Cecco M, Ito T, Petrashen AP, Elias AE, Skvir NJ, Criscione SW, Caligiana A, Brocculi G, Adney EM, Boeke JD et al. 2019. L1 drives IFN in senescent cells and promotes age-associated inflammation. Nature 566: 73–78.

Della Valle F, Reddy P, Yamamoto M, Liu P, Saera-Vila A, Bensaddek D, Zhang H, Prieto Martinez J, Abassi L, Celii M et al. 2022. LINE-1 RNA causes heterochromatin erosion and is a target for amelioration of senescent phenotypes in progeroid syndromes. Sci Transl Med 14: eabl6057.

Dobin A, Davis CA, Schlesinger F, Drenkow J, Zaleski C, Jha S, Batut P, Chaisson M, Gingeras TR. 2013. STAR: ultrafast universal RNA-seq aligner. *Bioinformatics (Oxford*, England) 29: 15–21.

Du M, Liu J, Chen X, Xie Y, Yuan C, Xiang Y, Sun B, Lan K, Chen M, James SJ et al. 2015. Casein kinase II controls TBK1/IRF3 activation in IFN response against viral infection. J Immunol 194: 4477–4488.

Eggenberger J, Blanco-Melo D, Panis M, Brennand KJ, tenOever BR. 2019. Type I interferon response impairs differentiation potential of pluripotent stem cells. Proc Natl Acad Sci U S A 116: 1384–1393.

Ernst J, Kellis M. 2012. ChromHMM: automating chromatin-state discovery and characterization. Nature Methods 9: 215–216.

Ernst J, Kellis M. 2017. Chromatin-state discovery and genome annotation with ChromHMM. Nat Protoc 12: 2478–2492.

Ewels PA, Peltzer A, Fillinger S, Patel H, Alneberg J, Wilm A, Garcia MU, Di Tommaso P, Nahnsen S. 2020. The nf-core framework for community-curated bioinformatics pipelines. Nature Biotechnology 38: 276–278.

Ewing AD, Kazazian HH, Jr. 2010. High-throughput sequencing reveals extensive variation in human-specific L1 content in individual human genomes. Genome research 20: 1262–1270.

Flemr M, Malik R, Franke V, Nejepinska J, Sedlacek R, Vlahovicek K, Svoboda P. 2013. A retrotransposon-driven dicer isoform directs endogenous small interfering RNA production in mouse oocytes. Cell 155: 807–816.

Gao D, Wu J, Wu YT, Du F, Aroh C, Yan N, Sun L, Chen ZJ. 2013. Cyclic GMP-AMP synthase is an innate immune sensor of HIV and other retroviruses. Science 341: 903–906.

Golding MC, Zhang L, Mann MR. 2010. Multiple epigenetic modifiers induce aggressive viral extinction in extraembryonic endoderm stem cells. Cell Stem Cell 6: 457–467.

Goodier JL, Cheung LE, Kazazian HH, Jr. 2012. MOV10 RNA helicase is a potent inhibitor of retrotransposition in cells. PLoS Genet 8: e1002941.

Goodier JL, Ostertag EM, Du K, Kazazian HH, Jr. 2001. A novel active L1 retrotransposon subfamily in the mouse. Genome research 11: 1677–1685.

Grant CE, Bailey TL, Noble WS. 2011. FIMO: scanning for occurrences of a given motif. Bioinformatics 27: 1017–1018.

Grow EJ, Flynn RA, Chavez SL, Bayless NL, Wossidlo M, Wesche DJ, Martin L, Ware CB, Blish CA, Chang HY et al. 2015. Intrinsic retroviral reactivation in human preimplantation embryos and pluripotent cells. Nature 522: 221–225.

Hamdorf M, Idica A, Zisoulis DG, Gamelin L, Martin C, Sanders KJ, Pedersen IM. 2015. miR-128 represses L1 retrotransposition by binding directly to L1 RNA. Nat Struct Mol Biol 22: 824–831.

Hertzog PJ, Hwang SY, Kola I. 1994. Role of interferons in the regulation of cell proliferation, differentiation, and development. Mol Reprod Dev 39: 226–232.

Herzner AM, Hagmann CA, Goldeck M, Wolter S, Kubler K, Wittmann S, Gramberg T, Andreeva L, Hopfner KP, Mertens C et al. 2015. Sequence-specific activation of the DNA sensor cGAS by Y-form DNA structures as found in primary HIV-1 cDNA. Nat Immunol 16: 1025–1033.

Iskow RC, McCabe MT, Mills RE, Torene S, Pittard WS, Neuwald AF, Van Meir EG, Vertino PM, Devine SE. 2010. Natural mutagenesis of human genomes by endogenous retrotransposons. Cell 141: 1253–1261.

Jachowicz JW, Bing X, Pontabry J, Boskovic A, Rando OJ, Torres-Padilla ME. 2017. LINE-1 activation after fertilization regulates global chromatin accessibility in the early mouse embryo. Nature genetics 49: 1502–1510.

Kano H, Godoy I, Courtney C, Vetter MR, Gerton GL, Ostertag EM, Kazazian HH, Jr. 2009. L1 retrotransposition occurs mainly in embryogenesis and creates somatic mosaicism. Genes & development 23: 1303–1312.

Kassiotis G, Stoye JP. 2016. Immune responses to endogenous retroelements: taking the bad with the good. Nat Rev Immunol 16: 207–219.

Kazazian HH, Jr., Moran JV. 2017. Mobile DNA in Health and Disease. N Engl J Med 377: 361–370.

Kazazian HH, Jr., Wong C, Youssoufian H, Scott AF, Phillips DG, Antonarakis SE. 1988. Haemophilia A resulting from de novo insertion of L1 sequences represents a novel mechanism for mutation in man. Nature 332: 164–166.

Krueger F. 2021. FelixKrueger/TrimGalore: v0.6.7. Zenodo.

Lee E, Iskow R, Yang L, Gokcumen O, Haseley P, Luquette LJ, 3rd, Lohr JG, Harris CC, Ding L, Wilson RK et al. 2012. Landscape of somatic retrotransposition in human cancers. Science 337: 967–971.

Leviyang S. 2021. Interferon stimulated binding of ISRE is cell type specific and is predicted by homeostatic chromatin state. Cytokine X 3: 100056.

Li H, Durbin R. 2009. Fast and accurate short read alignment with Burrows–Wheeler transform. Bioinformatics 25: 1754–1760.

Lin R, Mamane Y, Hiscott J. 1999. Structural and functional analysis of interferon regulatory factor 3: localization of the transactivation and autoinhibitory domains. Mol Cell Biol 19: 2465–2474.

Liu X, Liu Z, Wu Z, Ren J, Fan Y, Sun L, Cao G, Niu Y, Zhang B, Ji Q et al. 2023. Resurrection of endogenous retroviruses during aging reinforces senescence. Cell 186: 287–304 e226.

Love MI, Huber W, Anders S. 2014. Moderated estimation of fold change and dispersion for RNA-seq data with DESeq2. Genome Biology 15: 550.

Luo WW, Lian H, Zhong B, Shu HB, Li S. 2016. Kruppel-like factor 4 negatively regulates cellular antiviral immune response. Cell Mol Immunol 13: 65–72.

Luo Y, Hitz BC, Gabdank I, Hilton JA, Kagda MS, Lam B, Myers Z, Sud P, Jou J, Lin K et al. 2020. New developments on the Encyclopedia of DNA Elements (ENCODE) data portal. Nucleic Acids Research 48: D882–D889.

Ma W, Huang G, Wang Z, Wang L, Gao Q. 2023. IRF7: role and regulation in immunity and autoimmunity. Front Immunol 14: 1236923.

Macfarlan TS, Gifford WD, Agarwal S, Driscoll S, Lettieri K, Wang J, Andrews SE, Franco L, Rosenfeld MG, Ren B et al. 2011. Endogenous retroviruses and neighboring genes are coordinately repressed by LSD1/KDM1A. Genes & development 25: 594–607.

Macfarlan TS, Gifford WD, Driscoll S, Lettieri K, Rowe HM, Bonanomi D, Firth A, Singer O, Trono D, Pfaff SL. 2012. Embryonic stem cell potency fluctuates with endogenous retrovirus activity. Nature 487: 57–63.

Maillard PV, Ciaudo C, Marchais A, Li Y, Jay F, Ding SW, Voinnet O. 2013. Antiviral RNA interference in mammalian cells. Science 342: 235–238.

Maillard PV, Van der Veen AG, Deddouche-Grass S, Rogers NC, Merits A, Reis e Sousa C. 2016. Inactivation of the type I interferon pathway reveals long double-stranded RNA-mediated RNA interference in mammalian cells. EMBO J 35: 2505–2518.

Malki S, van der Heijden GW, O’Donnell KA, Martin SL, Bortvin A. 2014. A role for retrotransposon LINE-1 in fetal oocyte attrition in mice. Dev Cell 29: 521–533.

Marasca F, Sinha S, Vadala R, Polimeni B, Ranzani V, Paraboschi EM, Burattin FV, Ghilotti M, Crosti M, Negri ML et al. 2022. LINE1 are spliced in non-canonical transcript variants to regulate T cell quiescence and exhaustion. Nature genetics 54: 180–193.

McLeay RC, Bailey TL. 2010. Motif Enrichment Analysis: a unified framework and an evaluation on ChIP data. BMC Bioinformatics 11: 165.

Miyoshi T, Makino T, Moran JV. 2019. Poly(ADP-Ribose) Polymerase 2 Recruits Replication Protein A to Sites of LINE-1 Integration to Facilitate Retrotransposition. Mol Cell 75: 1286–1298 e1212.

Modzelewski AJ, Shao W, Chen J, Lee A, Qi X, Noon M, Tjokro K, Sales G, Biton A, Anand A et al. 2021. A mouse-specific retrotransposon drives a conserved Cdk2ap1 isoform essential for development. Cell 184: 5541–5558 e5522.

Moran JV, Holmes SE, Naas TP, DeBerardinis RJ, Boeke JD, Kazazian HH, Jr. 1996. High frequency retrotransposition in cultured mammalian cells. Cell 87: 917–927.

Muckenhuber M, Seufert I, Muller-Ott K, Mallm JP, Klett LC, Knotz C, Hechler J, Kepper N, Erdel F, Rippe K. 2023. Epigenetic signals that direct cell type-specific interferon beta response in mouse cells. Life Sci Alliance 6.

Muotri AR, Chu VT, Marchetto MC, Deng W, Moran JV, Gage FH. 2005. Somatic mosaicism in neuronal precursor cells mediated by L1 retrotransposition. Nature 435: 903–910.

Newkirk SJ, Lee S, Grandi FC, Gaysinskaya V, Rosser JM, Vanden Berg N, Hogarth CA, Marchetto MCN, Muotri AR, Griswold MD et al. 2017. Intact piRNA pathway prevents L1 mobilization in male meiosis. Proc Natl Acad Sci U S A 114: E5635–E5644.

Nielsen MI, Wolters JC, Bringas OGR, Jiang H, Di Stefano LH, Oghbaie M, Hozeifi S, Nitert MJ, van Pijkeren A, Smit M et al. 2025. Targeted detection of endogenous LINE-1 proteins and ORF2p interactions. Mob DNA 16: 3.

Ogata JD, Mu W, Davis ES, Xue B, Harrell JC, Sheffield NC, Phanstiel DH, Love MI, Dozmorov MG. 2023. excluderanges: exclusion sets for T2T-CHM13, GRCm39, and other genome assemblies. Bioinformatics 39: btad198.

Ohno R, Nakayama M, Naruse C, Okashita N, Takano O, Tachibana M, Asano M, Saitou M, Seki Y. 2013. A replication-dependent passive mechanism modulates DNA demethylation in mouse primordial germ cells. Development 140: 2892–2903.

Ozata DM, Gainetdinov I, Zoch A, O’Carroll D, Zamore PD. 2019. PIWI-interacting RNAs: small RNAs with big functions. Nat Rev Genet 20: 89–108.

Paludan SR, Bowie AG. 2013. Immune sensing of DNA. Immunity 38: 870–880.

Patel H, Espinosa-Carrasco J, Langer B, Ewels P, bot n-c, Garcia MU, Syme R, Peltzer A, Talbot A, Behrens D et al. 2023. nf-core/atacseq: [2.1.2] - 2022-08-07. Zenodo.

Patel H, Espinosa-Carrasco J, Wang C, Ewels P, bot n-c, Silva TC, Peltzer A, Langer B, Guinchard S, Garcia MU et al. 2024a. nf-core/chipseq: nf-core/chipseq v2.1.0 - Platinum Willow Sparrow. Zenodo.

Patel H, Ewels P, Manning J, Garcia MU, Peltzer A, Hammarén R, Botvinnik O, Talbot A, Sturm G, bot n-c et al. 2024b. nf-core/rnaseq: nf-core/rnaseq v3.18.0 - Lithium Lynx. Zenodo.

Patel S, Blaauboer SM, Tucker HR, Mansouri S, Ruiz-Moreno JS, Hamann L, Schumann RR, Opitz B, Jin L. 2017. The Common R71H-G230A-R293Q Human TMEM173 Is a Null Allele. J Immunol 198: 776–787.

Patro R, Duggal G, Love MI, Irizarry RA, Kingsford C. 2017. Salmon provides fast and bias-aware quantification of transcript expression. Nature Methods 14: 417–419.

Percharde M, Lin C-J, Yin Y, Guan J, Peixoto GA, Bulut-Karslioglu A, Biechele S, Huang B, Shen X, Ramalho-Santos M. 2018. A LINE1-nucleolin partnership regulates early development and ESC identity. Cell 174: 391–405. e319.

Poirier EZ, Buck MD, Chakravarty P, Carvalho J, Frederico B, Cardoso A, Healy L, Ulferts R, Beale R, Reis e Sousa C. 2021. An isoform of Dicer protects mammalian stem cells against multiple RNA viruses. Science 373: 231–236.

Ramírez F, Ryan DP, Grüning B, Bhardwaj V, Kilpert F, Richter AS, Heyne S, Dündar F, Manke T. 2016. deepTools2: a next generation web server for deep-sequencing data analysis. Nucleic Acids Research 44: W160–165.

Richardson SR, Gerdes P, Gerhardt DJ, Sanchez-Luque FJ, Bodea GO, Munoz-Lopez M, Jesuadian JS, Kempen MHC, Carreira PE, Jeddeloh JA et al. 2017. Heritable L1 retrotransposition in the mouse primordial germline and early embryo. Genome research 27: 1395–1405.

Risso D, Ngai J, Speed TP, Dudoit S. 2014. Normalization of RNA-seq data using factor analysis of control genes or samples. Nature Biotechnology 32: 896–902.

Robinson JT, Thorvaldsdóttir H, Winckler W, Guttman M, Lander ES, Getz G, Mesirov JP. 2011. Integrative Genomics Viewer. Nature biotechnology 29: 24–26.

Rodic N, Sharma R, Sharma R, Zampella J, Dai L, Taylor MS, Hruban RH, Iacobuzio-Donahue CA, Maitra A, Torbenson MS et al. 2014. Long interspersed element-1 protein expression is a hallmark of many human cancers. Am J Pathol 184: 1280–1286.

Rowland BD, Bernards R, Peeper DS. 2005. The KLF4 tumour suppressor is a transcriptional repressor of p53 that acts as a context-dependent oncogene. Nat Cell Biol 7: 1074–1082.

Sasaki N, Homme M, Kitajima S. 2023. Targeting the loss of cGAS/STING signaling in cancer. Cancer Sci 114: 3806–3815.

Schorn AJ, Gutbrod MJ, LeBlanc C, Martienssen R. 2017. LTR-Retrotransposon Control by tRNA-Derived Small RNAs. Cell 170: 61–71 e11.

Scopa C, Barnada SM, Cicardi ME, Singer M, Trotti D, Trizzino M. 2023. JUN upregulation drives aberrant transposable element mobilization, associated innate immune response, and impaired neurogenesis in Alzheimer’s disease. Nat Commun 14: 8021.

Seisenberger S, Andrews S, Krueger F, Arand J, Walter J, Santos F, Popp C, Thienpont B, Dean W, Reik W. 2012. The dynamics of genome-wide DNA methylation reprogramming in mouse primordial germ cells. Mol Cell 48: 849–862.

Sethi A, Gu M, Gumusgoz E, Chan L, Yan KK, Rozowsky J, Barozzi I, Afzal V, Akiyama JA, Plajzer-Frick I et al. 2020. Supervised enhancer prediction with epigenetic pattern recognition and targeted validation. Nat Methods 17: 807–814.

Shukla R, Upton KR, Munoz-Lopez M, Gerhardt DJ, Fisher ME, Nguyen T, Brennan PM, Baillie JK, Collino A, Ghisletti S et al. 2013. Endogenous retrotransposition activates oncogenic pathways in hepatocellular carcinoma. Cell 153: 101–111.

Simon M, Van Meter M, Ablaeva J, Ke Z, Gonzalez RS, Taguchi T, De Cecco M, Leonova KI, Kogan V, Helfand SL et al. 2019. LINE1 Derepression in Aged Wild-Type and SIRT6-Deficient Mice Drives Inflammation. Cell Metab 29: 871–885 e875.

Sundaram V, Cheng Y, Ma Z, Li D, Xing X, Edge P, Snyder MP, Wang T. 2014. Widespread contribution of transposable elements to the innovation of gene regulatory networks. Genome research 24: 1963–1976.

Sundaram V, Wang T. 2018. Transposable Element Mediated Innovation in Gene Regulatory Landscapes of Cells: Re-Visiting the “Gene-Battery” Model. Bioessays 40.

Tam OH, Aravin AA, Stein P, Girard A, Murchison EP, Cheloufi S, Hodges E, Anger M, Sachidanandam R, Schultz RM et al. 2008. Pseudogene-derived small interfering RNAs regulate gene expression in mouse oocytes. Nature 453: 534–538.

Thomas CA, Tejwani L, Trujillo CA, Negraes PD, Herai RH, Mesci P, Macia A, Crow YJ, Muotri AR. 2017. Modeling of TREX1-Dependent Autoimmune Disease using Human Stem Cells Highlights L1 Accumulation as a Source of Neuroinflammation. Cell Stem Cell 21: 319–331 e318.

Torres J, Watt FM. 2008. Nanog maintains pluripotency of mouse embryonic stem cells by inhibiting NFkappaB and cooperating with Stat3. Nat Cell Biol 10: 194–201.

Tristan-Ramos P, Rubio-Roldan A, Peris G, Sanchez L, Amador-Cubero S, Viollet S, Cristofari G, Heras SR. 2020. The tumor suppressor microRNA let-7 inhibits human LINE-1 retrotransposition. Nat Commun 11: 5712.

Vorontsov IE, Eliseeva IA, Zinkevich A, Nikonov M, Abramov S, Boytsov A, Kamenets V, Kasianova A, Kolmykov S, Yevshin Ivan S et al. 2024. HOCOMOCO in 2024: a rebuild of the curated collection of binding models for human and mouse transcription factors. Nucleic Acids Research 52: D154–D163.

Wang JT, Chang LS, Chen CJ, Doong SL, Chang CW, Chen MR. 2014a. Glycogen synthase kinase 3 negatively regulates IFN regulatory factor 3 transactivation through phosphorylation at its linker region. Innate Immun 20: 78–87.

Wang R, Teng C, Spangler J, Wang J, Huang F, Guo YL. 2014b. Mouse embryonic stem cells have underdeveloped antiviral mechanisms that can be exploited for the development of mRNA-mediated gene expression strategy. Stem Cells Dev 23: 594–604.

Wang R, Wang J, Acharya D, Paul AM, Bai F, Huang F, Guo YL. 2014c. Antiviral responses in mouse embryonic stem cells: differential development of cellular mechanisms in type I interferon production and response. J Biol Chem 289: 25186–25198.

Wang R, Wang J, Paul AM, Acharya D, Bai F, Huang F, Guo YL. 2013. Mouse embryonic stem cells are deficient in type I interferon expression in response to viral infections and double-stranded RNA. J Biol Chem 288: 15926–15936.

Witteveldt J, Knol LI, Macias S. 2019. MicroRNA-deficient mouse embryonic stem cells acquire a functional interferon response. Elife 8.

Witteveldt J, Liu Z, Ariza-Cosano A, Ramirez C, Walters JL, Marchante PG, Maas L, Ivens A, Tebaldi T, Heras SR et al. 2025. Double stranded RNA sensing drives interferon silencing in early development. bioRxiv: 2025.2003.2010.642424.

Wu T, Hu E, Xu S, Chen M, Guo P, Dai Z, Feng T, Zhou L, Tang W, Zhan L et al. 2021. clusterProfiler 4.0: A universal enrichment tool for interpreting omics data. The Innovation 2: 100141.

Wu X, Dao Thi VL, Huang Y, Billerbeck E, Saha D, Hoffmann HH, Wang Y, Silva LAV, Sarbanes S, Sun T et al. 2018. Intrinsic Immunity Shapes Viral Resistance of Stem Cells. Cell 172: 423–438 e425.

Yockey LJ, Iwasaki A. 2018. Interferons and Proinflammatory Cytokines in Pregnancy and Fetal Development. Immunity 49: 397–412.

Yu C, Wang B, Zhu Y, Zhang C, Ren L, Lei X, Xiang Z, Zhou Z, Huang H, Wang J et al. 2022. ID2 inhibits innate antiviral immunity by blocking TBK1- and IKKepsilon-induced activation of IRF3. Sci Signal 15: eabh0068.

Yu G. 2025. enrichplot: Visualization of Functional Enrichment Result.

Yue F Cheng Y Breschi A Vierstral J Wu W Ryba T Sandstrom R Ma Z Davis C Pope BD et al. 2014. A comparative encyclopedia of DNA elements in the mouse genome. Nature 515: 355–364.

Zhang H, Li J, Yu Y, Ren J, Liu Q, Bao Z, Sun S, Liu X, Ma S, Liu Z et al. 2023. Nuclear lamina erosion-induced resurrection of endogenous retroviruses underlies neuronal aging. Cell Rep 42: 113396.

Zhao Y, Oreskovic E, Zhang Q, Lu Q, Gilman A, Lin YS, He J, Zheng Z, Lu JY, Lee J et al. 2021. Transposon-triggered innate immune response confers cancer resistance to the blind mole rat. Nat Immunol 22: 1219–1230.

Zoch A, Konieczny G, Auchynnikava T, Stallmeyer B, Rotte N, Heep M, Berrens RV, Schito M, Kabayama Y, Schopp T et al. 2024. C19ORF84 connects piRNA and DNA methylation machineries to defend the mammalian germ line. Mol Cell 84: 1021–1035 e1011.

