## Supplementary material for "Inhibition of cytosolic DNA sensing and transposon activity safeguards pluripotency": Supp figures

**List of data:**

Figure S1
Figure S2
Figure S3
Figure S4
Figure S5
Figure S6

Table S1: normalized RNA-seq expression data
Table S2: Published ATAC/ChIP-seq analysis and ChromHMM model generation
Table S3: list of primers, sgRNAs and antibodies

### Figure S1

Garcia-Llagostera et al.

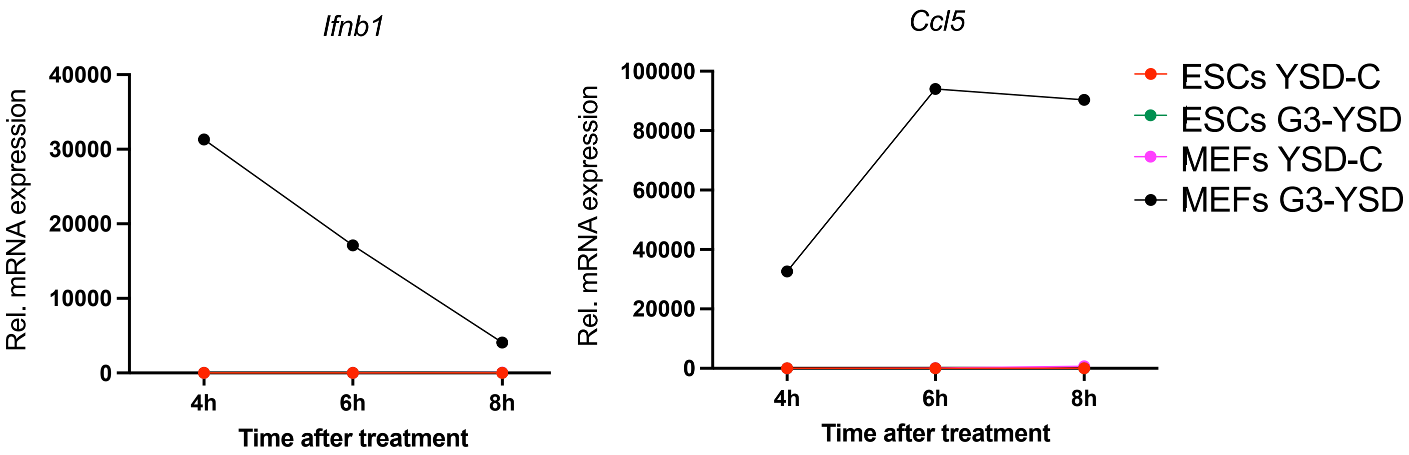


**Figure S1. Response of ESCs and MEFs to cGAS/STING stimulation**
RT-qPCR analysis of the selected genes following a timecourse after transfection of G3-YSD or YSD-C ligands. 4h YSD DNA control in ESCs is set to 1.

**Figure S2** **Garcia-Llagostera et al.**

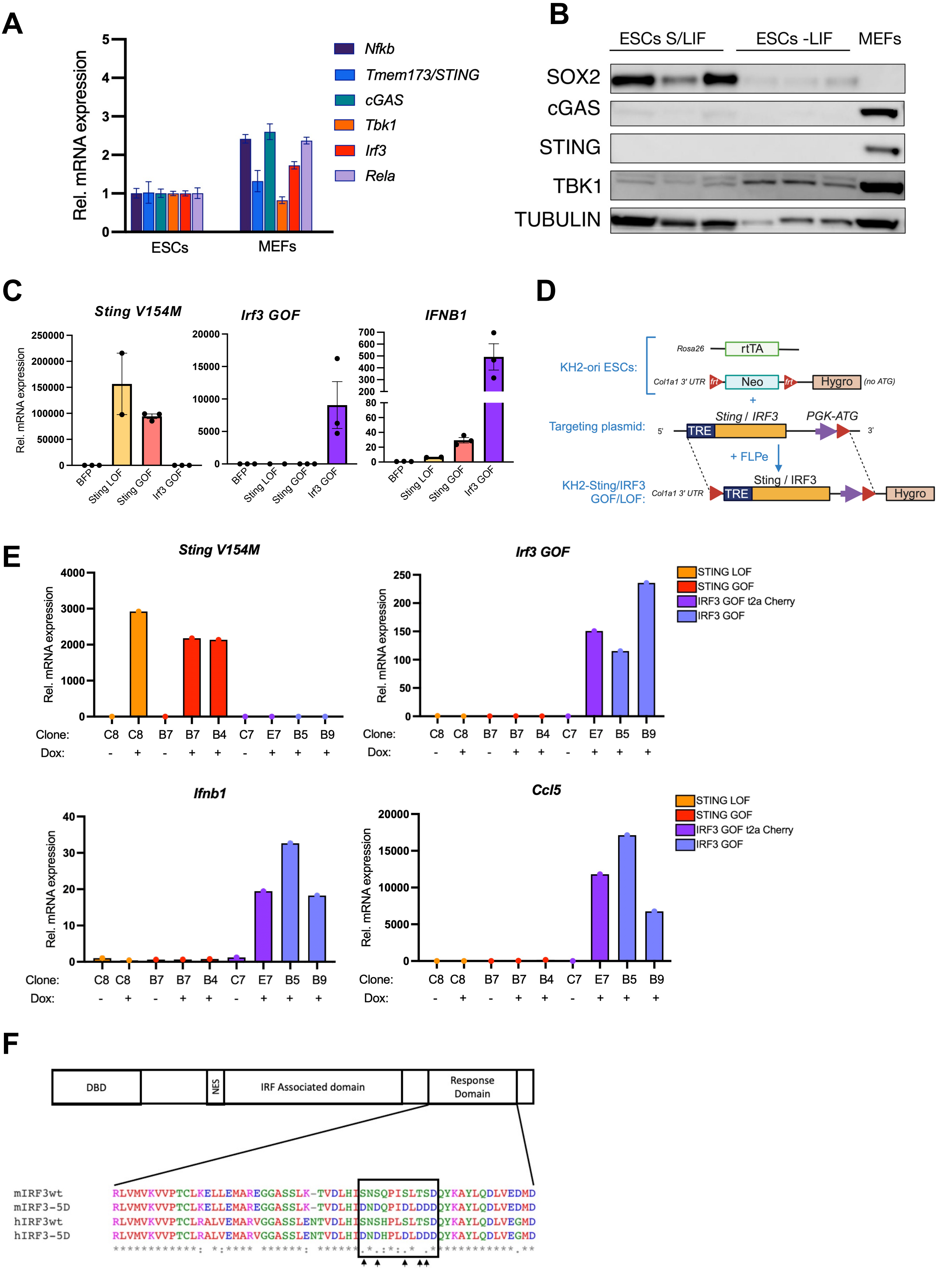

**Figure S2. cGAS/STING expression in ESCs and MEFs**

A) RT-qPCR analysis of genes involved in the cGAS/STING pathway in ESCs and MEFs. Data are mean +/- n=3, representative of 3 experiments.

B) Western blot in ESCs and MEFs showing the absence of cGAS and STING protein in ESCs and upon differentiation (samples from n=3 experiments). TUBULIN is shown as a loading control.

C) RT-qPCR analysis of construct over-expression (*Sting/Irf3*) or human *IFNB* expression in 293T cells following transient transfection of the indicated constructs in the presence of Dox. Sting-GOF and Irf3-GOF induce *IFNB* expression but not Sting-LOF. Data are shown plus mean +/- s.e.m of 2-3 wells, representative of 2 independent experiments

D) Generation of *Sting/Irf3* stable cell lines. The Parental ESC line (KH2-ori) contains an frt-flanked Neomycin resistance cassette, followed by a Hygromycin resistance cassette that is missing an ATG start codon. Separately, they constitutively express the rtTA transactivator protein from the *Rosa26* locus. Flippase (FLPe) mediated recombination of a *Sting/IRF* expression plasmid replaces the Neo cassette with the Dox-inducible construct, as well as a PGK promoter and ATG start codon upstream of Hygro. The resultant colonies are HygroR and express the construct of interest under the control of Doxycycline.

E) RT-qPCR analysis of gene expression in stable KH2 Dox-inducible cell lines. For IRF3 GOF, clones with or without downstream t2a-mCherry expression were generated and analyzed. Experiments were performed with multiple clones per construct and are representative of 2 experiments. Subsequent IRF3 GOF experiments were performed with E7, B5 and/or B9.

F) Alignment of IRF3 protein sequences from mouse and human, with arrows indicating the analogous Ser/Thr residues in mouse that have been mutated to aspartate (D) for IRF3 GOF and constitutive activity.

**Figure S3** **Garcia-Llagostera et al.**

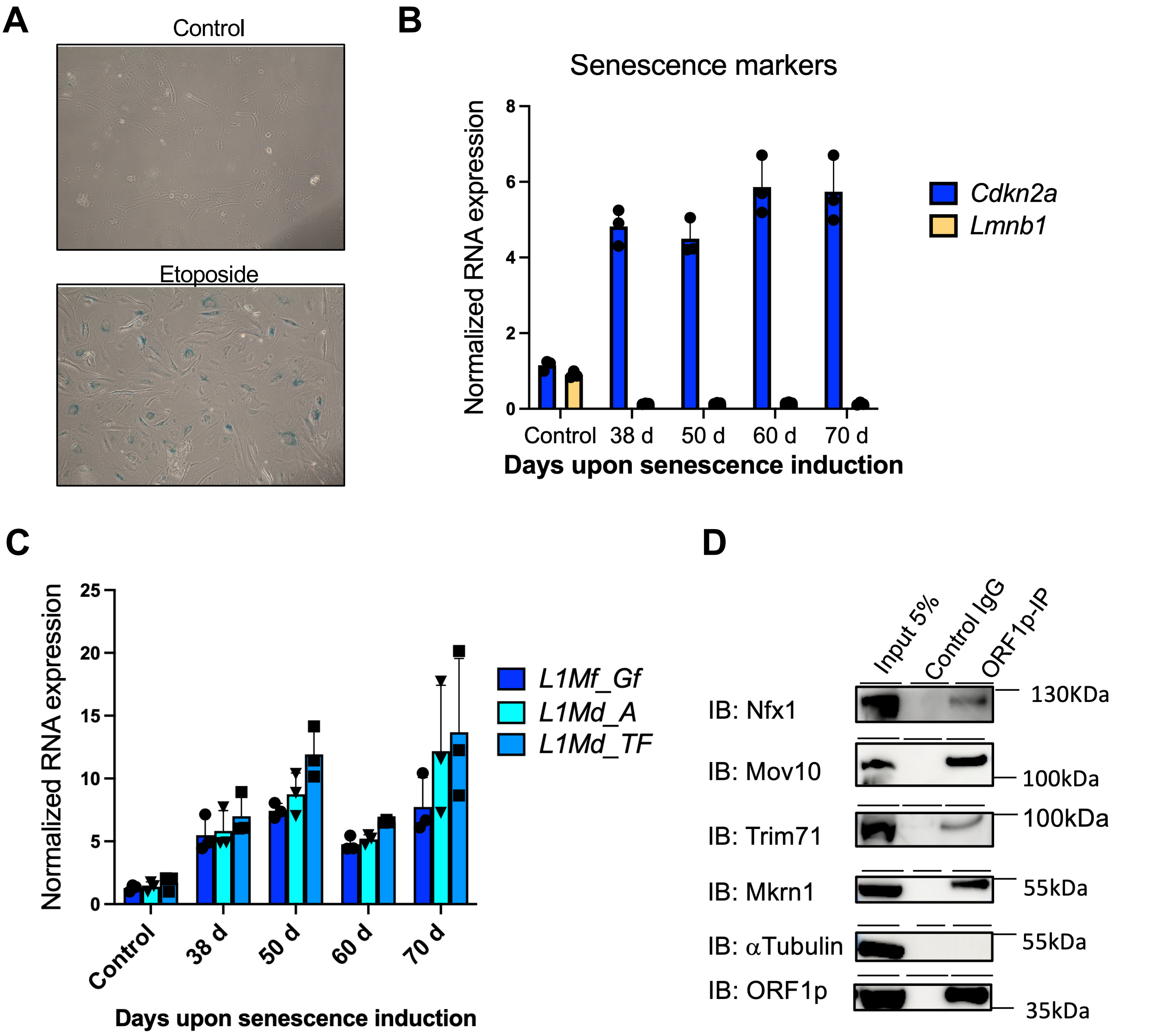


**Figure S3. Post-transcriptional repression of LINE1 activity in ESCs**

a) Staining of control and senescent MEFs for β-galactosidase, 3 days after treatment with 20 μM Etoposide. MEFs were subsequently cultured further for the indicated number of days (following panels).
b) RT-qPCR analysis of senescence marker induction in MEFs after etoposide-induced senescence, cultured for the indicated time points. Data are mean +/- s.e.m, n=3 wells.
c) RT-qPCR analysis of young, transcriptionally active LINE1 subfamilies (*L1_Md*) expression in MEFs treated as in b).
d) Co-IP of novelly-identified LINE1 ORF1p interactors in ESCs. TUBULIN is shown as a negative control, and MOV10 as a positive control. Data in a-d are representative of at least 2 independent experiments.

### Figure S4 Garcia-Llagostera et al.

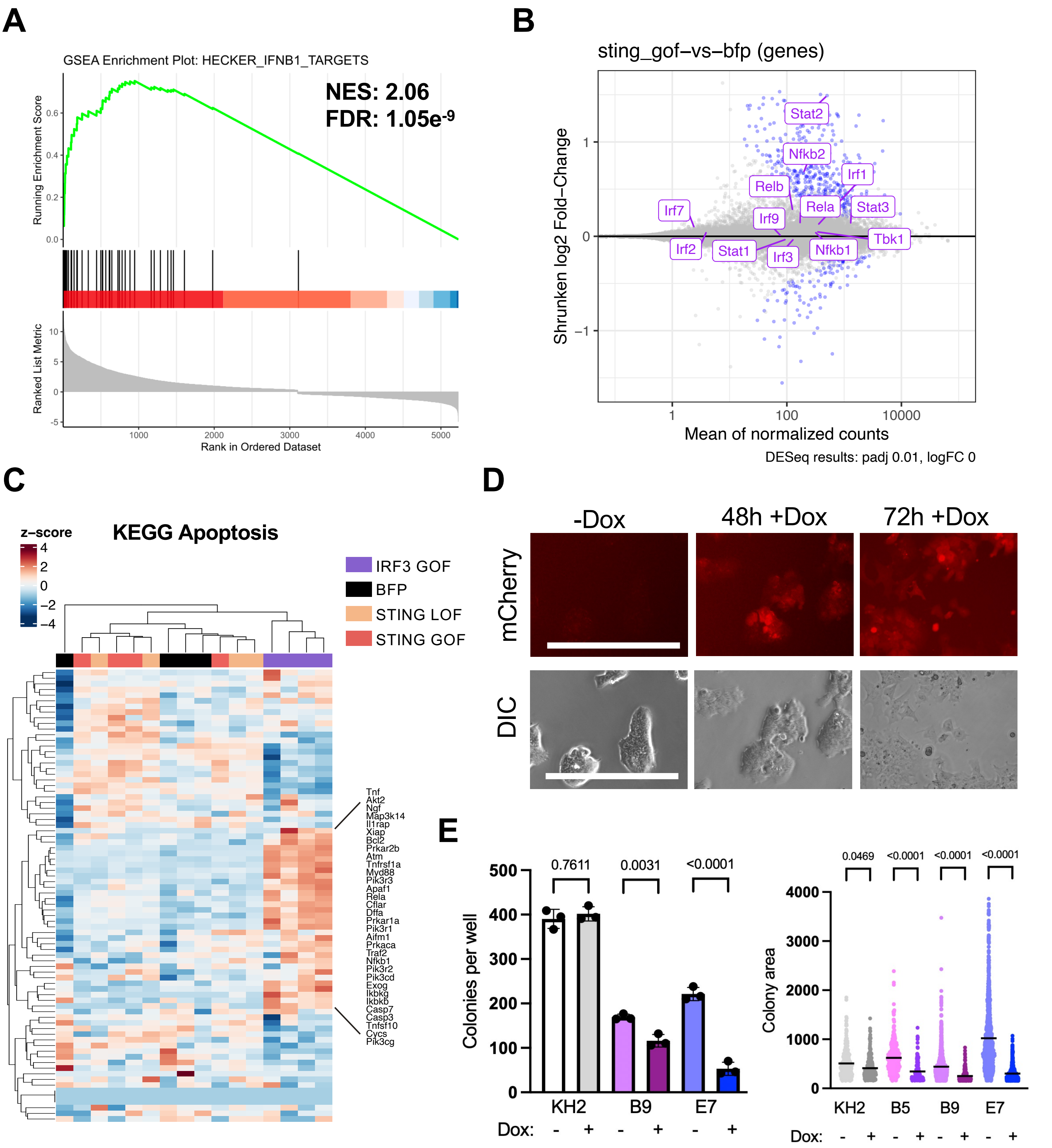


**Figure S4. IRF3 activation is incompatible with pluripotency**

A) GSEA analysis of IFN-I genes shows a significantly enrichment upon IRF3 GOF expression in ESCs.

B) MA plot showing $\log_2$FC in expression of genes in STING GOF-transfected versus BFP-transfected ESCs, highlighting selected IFN-I genes. Blue points indicate significant at FDR <0.01.

C) Hierarchically-clustered heatmap displaying normalized expression of KEGG apoptosis genes across conditions in ESCs.

D) Cherry expression is maintained in IRF3 GOF-t2a-cherry (E7) ESCs for at least 72h. Dox was refreshed daily. Scale bar, 300µm.

E) Colony formation assay in unmodified (KH2-ori) or IRF3 GOF ESCs, with and without the presence of Dox. Shown is one experiment, representative of 2-3 experiments. P values, One-way ANOVA with Sidak correction for multiple comparisons.

### Figure S5

Garcia-Llagostera et al.

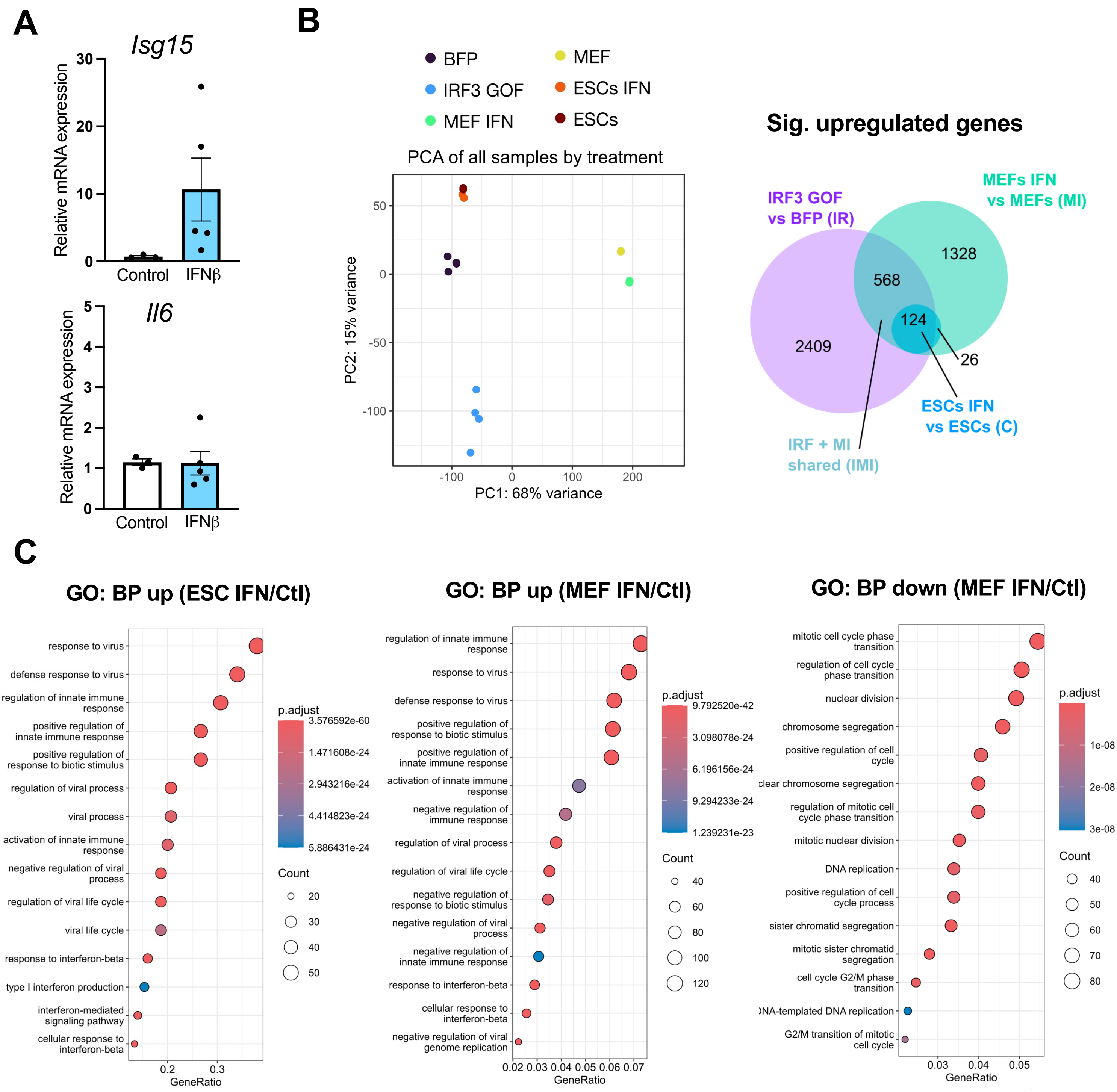


**Figure S5. IFN response in ESC and MEFs**

A) RT-qPCR of the indicated genes in blastocysts cultured for 6 h with or without the addition of 1000IU IFNβ. Data show n=3-5 independent batches of blastocysts treated independently with IFNβ.

B) PCA of normalized gene expression (vst) clusters samples by cell type along PC1 (68% variance) and treatment along PC2 (15% variance).

C) Enrichment of the top 15 significant biological processes (FDR < 0.05) identified in GO analysis of upregulated DEGs in the indicated comparisons.

### Figure S6

Garcia-Llagostera et al.

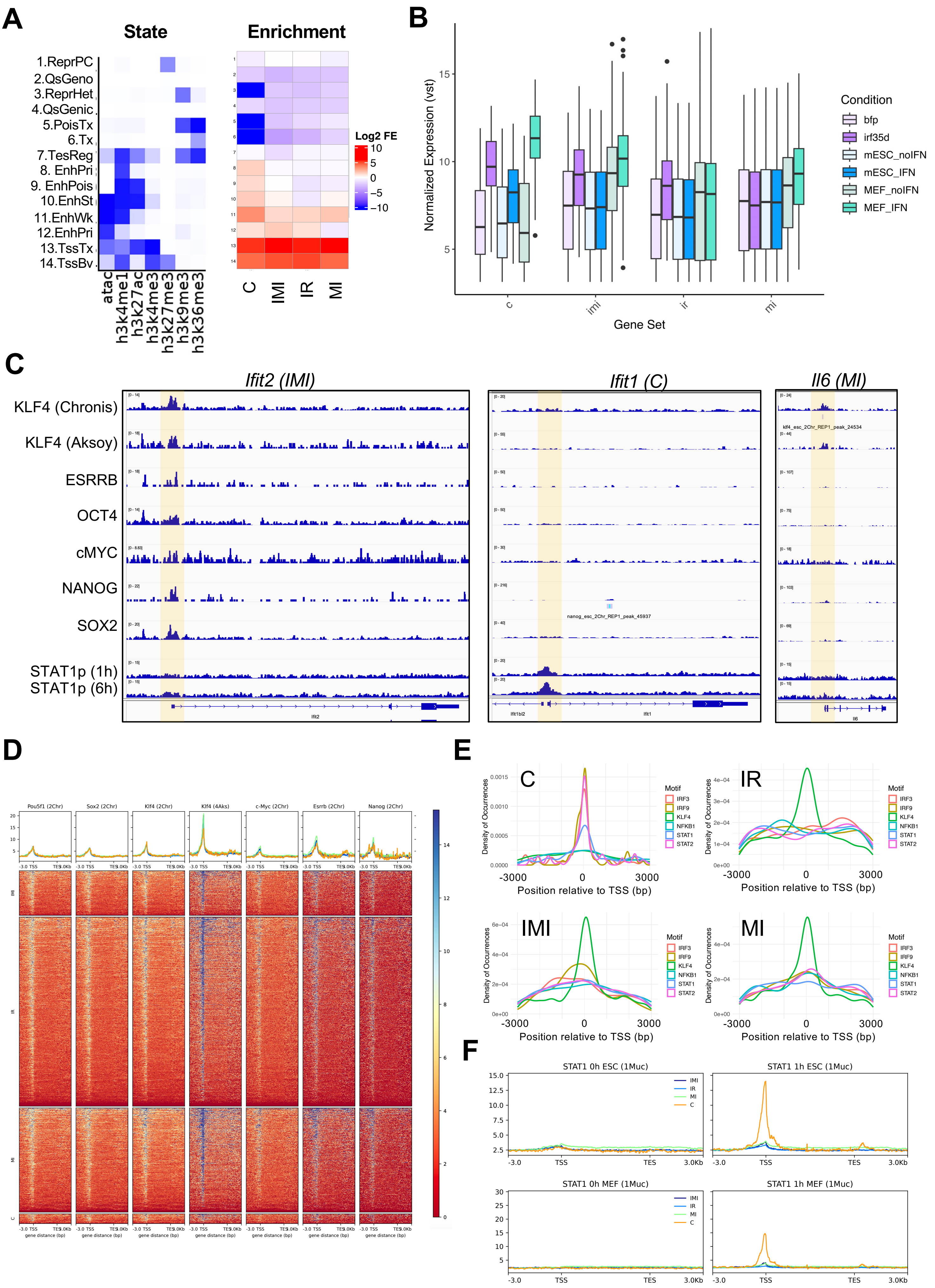

**Figure S6. IFN-I antagonism by the pluripotency network**

A) ChromHMM analysis using published ChIP-seq (Chronis et al., 2017, PMID: 28111071; Yue et al., 2014, PMID: 25409824; Sethi et al., 2020, PMID: 32737473) and ATAC-seq data (Muckenhuber et al., 2023, PMID: 36732019) in ESCs. Chromatin state emissions are displayed on the left, and enrichment of each state within the indicated gene sets is shown on the right. Further information and metrics supporting chromatin state annotations are provided in Table S2.

B) Grouped boxplot displaying vst-normalized expression of the indicated gene sets (C/IR/IMI/MI) across the different cell types and treatments.

C) ChIP enrichment tracks for the indicated transcription factors at examples from different classes of IFN-I genes. The gene sets (C/IR/IMI/MI) containing each gene are indicated in parentheses. ChIP-seq data are analysed from (Chronis et al., 2017), with an additional KLF4 dataset from (Aksoy et al., 2014, PMID: 24770696), and STAT1p from (Muckenhuber et al., 2023). The TSS is highlighted in yellow.

D) Spatial heatmaps of transcription factor binding across the indicated gene sets (C/IR/IMI/MI) – source datasets are as described in C.

E) Enrichment of the indicated transcription factor motifs ($P < 1e-5$) occurring +/- 3 Kb around the TSS of the C/IR/IMI/MI gene sets. Results are filtered for same-sense orientation relative to each gene.

F) STAT1p enrichment profiles (RPGC) over gene bodies (+/- 3kb) of each gene set (C/IR/IMI/MI) in IFNβ treated and untreated ESCs and MEFs. Data taken from (Muckenhuber et al., 2023).
